# Adaptation to bile and anaerobicity limits *Vibrio cholerae* phage adsorption

**DOI:** 10.1101/2023.04.22.537938

**Authors:** Zoe Netter, Drew T. Dunham, Kimberley D. Seed

## Abstract

Bacteriophages (viruses of bacteria) play a pivotal role in shaping both the evolution and dynamics of bacterial populations. Bacteria employ arsenals of genetically encoded phage defense systems, but can alternatively achieve protection by changing the availability of cellular resources that phages rely on for propagation. These physiological changes are often adaptive responses to unique environmental signals. The facultative pathogen *Vibrio cholerae* adapts to both aquatic and intestinal environments with niche-specific physiological changes that ensure its evolutionary success in such disparate settings. In both niches, *V. cholerae* is susceptible to predation by lytic phages like ICP1. However, both phages and susceptible bacterial hosts coexist in nature, indicating that environmental cues may modulate *V. cholerae* cell state to protect against phage infection. This work explores one such modification in response to the intestine-specific signals of bile and anaerobicity. We found that *V. cholerae* grown in these conditions reduces O1-antigen decoration on its outer membrane lipopolysaccharide. Because the O1-antigen is an essential moiety for ICP1 phage infection, we investigated the effect of partial O1-antigen depletion as a mechanism of phage defense and observed that O1-depletion limits phage adsorption. We identified mechanistic contributions to O1-depletion, including the essentiality of a weak acid tolerance system for O1 production at low pH, and alterations in transcriptional profiles indicating limitations in resources for O1-biosynthesis. This analysis illustrates a complex interplay between signals relevant to the intestinal environment and bacterial physiology that provide *V. cholerae* with protection from phage predation.

## Introduction

Bacteria thrive in spite of the constant threat of infection by predatory bacteriophages (or phages, viruses of bacteria). Selective pressure imposed by phage predation shapes bacterial populations and evolutionary trajectories. As much as 10% of a bacterial genome can be dedicated to phage defense^2^, and phage defense genes can constitute major genomic differences contributing to inter- and intra-species diversity^3^. There has been a recent acceleration in the discovery of diverse and innovative systems that defend bacteria from phage predation, from just a few simple genes providing defense like restriction-modification systems^4^ to complex systems like CRISPR-Cas^5^ and phage satellites^6, 7^. However, a bacterium’s capacity to limit phage infection does not depend only on genetically encoded systems. Bacterial physiology also plays a pivotal role in phage defense. Global bacterial transcription and translation activity, metabolism, and availability of cellular components can impact the efficiency of phage infection, even in nearly identical host strains^8^. While some phage defense systems have been shown to manipulate bacterial physiology to favor host population survival (such as a group of Sir2-domain-containing systems that respond to phage infection by degrading cellular NAD+, a necessary coenzyme in central carbon metabolism^9–11)^, the impact of bacterial host physiology on phage infection is largely not mechanistically understood. This knowledge gap is compounded by the dynamic interplay between cell state and environment, which is fairly static in a laboratory setting but varies dramatically in a bacterium’s natural environment.

*Vibrio cholerae*, the bacterial agent that causes the diarrheal disease cholera, has widely variable transcriptional activity and metabolic states in its natural ecological niches. *V. cholerae* has evolved several intricate mechanisms to sense its environment and alter cellular state accordingly to adapt rapidly to disparate conditions. When the bacteria are ingested from an aquatic reservoir and enter a human host, they sense environmental cues like bile acids, anaerobicity, pH, and changes in osmolarity and temperature^12, 13^. These intestinal stimuli trigger the *V. cholerae* ToxR regulon virulence cascade, which (via ToxT) activates the expression of cholera toxin (CTX), toxin co-regulated pilus (TCP), and other factors necessary to establish infection and support cholera pathogenesis^12^. Late in infection, gene expression shifts to make global alterations in metabolism in preparation for persistence in the stool or aquatic reservoir^14^, where a different set of genes appears necessary for survival^15^. In addition to fluctuating environmental conditions, *V. cholerae* also encounters predation by lytic phages in both the human gut and the aquatic reservoir^16, 17^. *V. cholerae* encodes a broad arsenal of anti-phage systems^7, 18, 19^, but beyond a few genetically encoded means to restrict phage infection, little is known about the influence of pertinent intestinal stimuli on *V. cholerae’s* susceptibility to phage predation. There is evidence that genetically “defenseless” strains can be co-isolated with lytic phages from cholera patient stool samples, indicating co-existence in the human gut^18^. However, it is unclear whether this is due to spatial separation resulting from the heterogeneity of the gut environment and/or by changes in bacterial physiology and metabolism in the gut environment that also impact phage susceptibility.

The bacterial cell surface provides the first line of defense against phage attack and environmental challenge. In Gram-negative bacteria like *V. cholerae*, the surface outer membrane is composed of a phospholipid bilayer embedded with lipopolysaccharide (LPS, composed of a bilayer-embedded lipid A component, a core-polysaccharide linker, and a widely variable O-antigen polysaccharide component) in the outer leaflet along with proteins involved in transmembrane transport and environmental sensing^20^. These outer membrane components are frequently recognized by phages as receptors during the attachment process^21, 22^. Phage receptor display can be modified directly by changes in the expression of their corresponding biosynthetic genes or by other changes in metabolism that alter the availability of the components necessary for biosynthesis or transport^20^. Although over 200 *V. cholerae* O-antigen serogroups have been identified, the vast majority of cholera infections are caused by the O1 serogroup^23^. The O1- antigen component of *V. cholerae* LPS is also the receptor for the dominant lytic vibriophage ICP1, which is recurrently co-isolated with *V. cholerae* in patient stool samples in areas where cholera is endemic^24^. *V. cholerae* is heavily restricted from permanently losing the LPS O1- antigen to escape ICP1 predation because it is required for efficient colonization of the intestine and subsequent pathogenesis^25^.

The coexistence of susceptible *V. cholerae* and ICP1 phages has been noted in several instances in cholera patient stool samples^16, 17, 26, 27^, leading us to hypothesize that some aspect of the human intestinal environment may alter phage susceptibility. Seeking to determine the physiological impact of specific gut-associated stimuli on *V. cholerae* and the repercussions for ICP1 infection, we focused specifically on bile acids and anaerobicity. Both of these stimuli are known to cause broad transcriptional changes in *V. cholerae*^13^. We assess the individual and combinatorial effects on *V. cholerae* LPS production and identify a combination of anaerobicity and bile acid supplementation that reduces the degree of O1-decoration of LPS in *V. cholerae*. We utilize phage-based assays to assess the biological relevance of this phenomenon and transcriptomics to identify several potential mechanisms for decreased O1-antigen production, including widespread transcriptional changes to central metabolic processes, reduced production of O1 biosynthetic enzymes and the essentiality of weak acid tolerance for O1-antigen production at low pH.

## Results

### *Vibrio cholerae* dynamically modifies the abundance of O1-decorated LPS in response to intestine-derived stimuli

To examine the effect of gut-specific cues on *V. cholerae* LPS, we cultured *V. cholerae* overnight in LB media supplemented with 0.5% bile both aerobically and anaerobically, then extracted and visualized LPS by denaturing gel electrophoresis and silver stain. *V. cholerae* cultured with bile supplementation alone or anaerobicity alone produced LPS similar to cells in standard LB aerobic culture, while *V. cholerae* grown in a combination of bile and anaerobic conditions produced LPS with strikingly less O1-decoration than the other conditions (Figure 1A). While the reduction in O1-antigen was visually apparent, it was not a complete depletion (∼30% less average O1 signal intensity) (Figure 1B).

**Figure 1.**
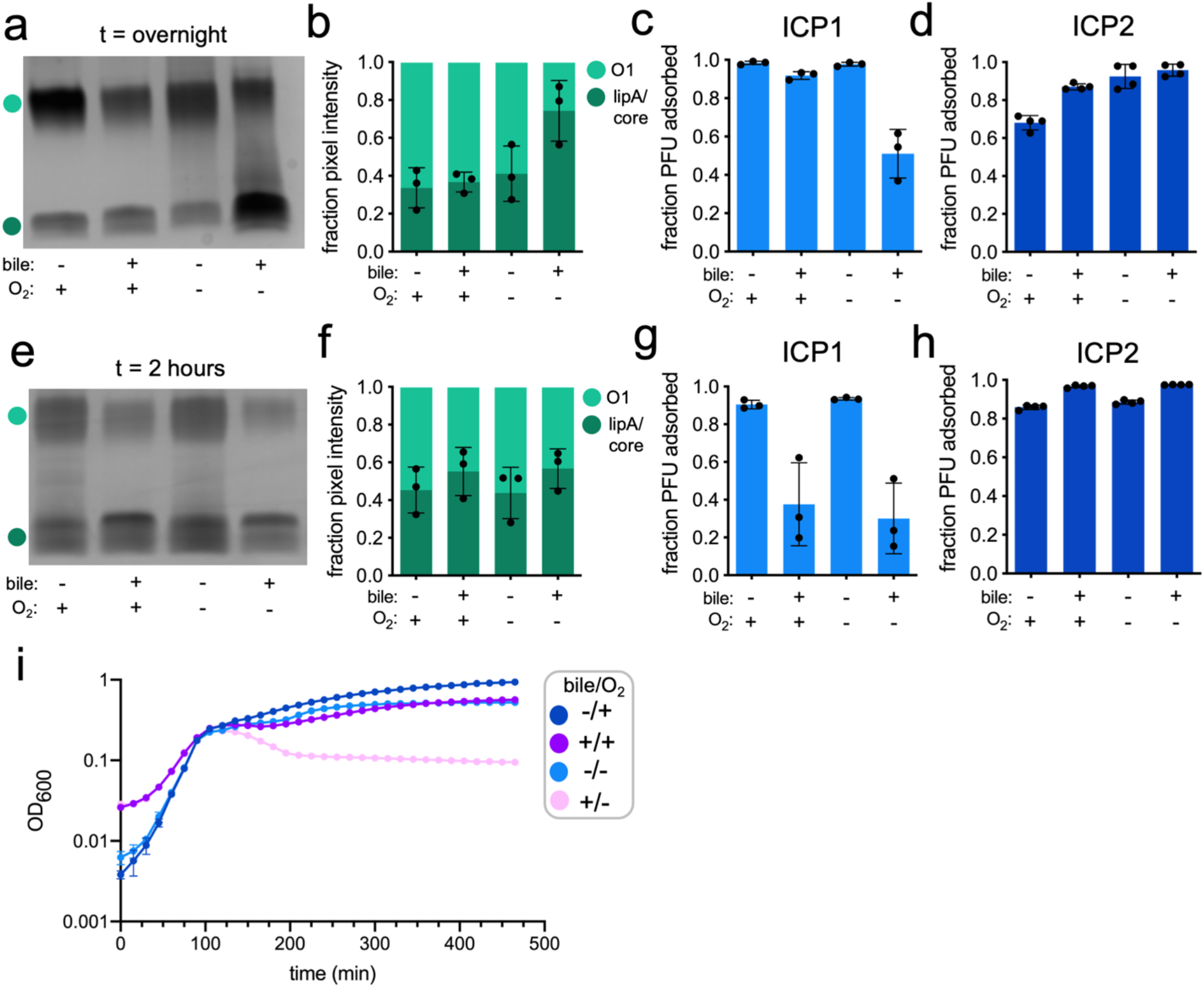
*Vibrio cholerae* dynamically modifies the abundance of O1-decorated LPS in response to intestine-derived stimuli (**a**) Silver stain of LPS purified from *V. cholerae* cultured overnight in LB or LB supplemented with 0.5% bile acids (bile -/+), aerobically or anaerobically (O_2_ +/-). (**b**) Average fraction of pixel intensity quantified from replicate purified LPS silver stains. Light green represents the relative fraction of O1-antigen signal and dark green represents the fraction of lipid A/core signal. (**c**) Fraction of O1-dependent phage ICP1 adsorbed by *V. cholerae* cultured overnight in all combinations of culture conditions. (**d**) Fraction of OmpU-dependent phage ICP2 adsorbed by *V. cholerae* cultured overnight in all combinations of culture conditions. (**e-h**) same as (a-d) with culture harvested at two hours. (**i**) Growth curve measuring OD_600_ for all combinations of culture conditions over time (in minutes).

We were curious to assess if the degree of reduction in O1-decorated LPS observed under anaerobic conditions with bile was biologically consequential for phage infection, specifically for the O1-dependent phage ICP1. We probed phage susceptibility in these different conditions with a modified adsorption assay, heat-killing cells grown in each condition to control for differences in host cell physiological state. The physiological state of host cells can dramatically alter the robustness of the phage life cycle after initial genome injection^28^, but measuring the capability of ICP1 to adsorb to heat-killed cells bypasses the challenges of comparing phage replication in hosts with different metabolic states and allows us to directly assess phage receptor availability. ICP1 efficiently adsorbed to *V. cholerae* cells cultured in standard, anaerobic, and aerobic bile conditions; however, ICP1 adsorption was reduced by ∼50% when cells were cultured anaerobically with bile (Figure 1C). Although the O1-antigen was not completely eliminated from LPS on cells in this condition, this indicates that the observed reduction in O1- decorated LPS has a sizeable impact on ICP1 adsorption. To further validate that the modified adsorption assay faithfully measures phage receptor availability, we repeated the experiment using ICP2, an unrelated vibriophage that uses the outer membrane porin OmpU as its receptor^29^. Expression of *ompU* is known to be upregulated in bile conditions^12, 30^. Concordantly, we observed enhanced ICP2 adsorption in bile-supplemented conditions (Figure 1D). These results validate that the reduction in O1-antigen abundance observed by purified LPS silver stain substantially impacts *V. cholerae* susceptibility to O1-dependent phages like ICP1, suggesting that such adaptations could affect *V. cholerae*’s susceptibility to phages during intestinal colonization.

Reduction in O1-decorated LPS can result from genetic mutations in the O1-antigen biosynthetic gene cluster^24^. If overnight growth in the anaerobic bile culture condition selects for mutants that produce less O1-antigen, then the reduced O1 phenotype from growth overnight in this condition would be expected to persist with outgrowth in the absence of such stimuli (i.e., in standard aerobic LB). In this case, O1 mutants would also be expected to have increased fitness in anaerobic bile conditions. Conversely, if the reduction in O1-decorated LPS is a result of transient phenotypic changes rather than permanent genetic changes, cultures grown overnight in anaerobic bile conditions would be expected to recover complete O1-antigen production when switched to standard conditions. To distinguish between these possibilities, we tested the recovery of anaerobic bile overnight *V. cholerae* cultures in standard aerobic conditions and observed complete recovery of O1-decorated LPS within 120 minutes (Supplementary Figure S1A-S1E). We also tested the growth efficiency of a genetic mutant (Δ*wbeL*) that produces no O1-antigen and found no difference in fitness relative to wild-type *V. cholerae* in either aerobic or anaerobic conditions with varying concentrations of bile (Supplementary Figure S1F, S1G).

We were also curious if this response was conserved among *V. cholerae* strains. We grew both classical *V. cholerae* and contemporary El Tor clinical isolates in anaerobic bile conditions and observed reductions in O1-antigen production similar to the *V. cholerae* E7946 strain used as our laboratory strain (which is also an El Tor clinical isolate) (Supplementary Figure S10), indicating that O1-antigen reduction is likely conserved across the serogroup. Taken together, these data suggest that the observed reduction in O1-decorated LPS produced by *V. cholerae* in anaerobic bile conditions is due to a reversible alteration in pathway(s) involved in O1-antigen biosynthesis.

LPS molecules are built from organic components of central carbon metabolism (the O1-antigen is synthesized from malate and D-fructose 6-phosphate)^31^. LPS biosynthesis may be impacted by the metabolic state of the cell simply due to different resource availability in different conditions. More stressful culture conditions could cause diminished production of central carbon metabolism components in order to decrease cell growth, as has been demonstrated in a dual transcriptomic and metabolomic analysis of the *E. coli* stress response^32^. Depletion of central carbon metabolism intermediates would effectively deplete the pool available for synthesizing the O1-antigen component of LPS, resulting in incomplete O1-decoration on the cell surface. We observed depletion of O1-decorated LPS in overnight cultures grown anaerobically with bile, a condition where cells may be actively altering their metabolic states to ensure continued survival.

To evaluate cell growth in all relevant conditions, we measured *V. cholerae* growth in each condition over time (Figure 1I). All cultures grew similarly for the first ∼120 minutes, however as the rest of the cultures proceeded to grow, the cells in bile anaerobic conditions instead persisted at a low but stable OD and CFU (Figure 1I and Supplementary Figure S9A). These data illustrate that while the end state of cells in each culture condition may vary dramatically, there are earlier instances where growth rates were more similar.

We were interested in assessing the state of cell surface LPS at the two-hour timepoint, where although the metabolic states of the cells are likely still fundamentally different, the relative growth rates appear similar regardless of culture condition. At two hours, we were surprised to observe a reduction in O1-decorated LPS in the presence of bile under both aerobic and anaerobic conditions. The difference was subtle by quantification of purified LPS silver stain (Figure 1E, 1F) but dramatic when measured by ICP1 adsorption assay (Figure 1G), while ICP2 adsorption remained fairly consistent across all conditions (Figure 1H). This suggested that, while the relative representation of O1 and lipid A/core components were difficult to ascertain by silver stain, the ICP1 adsorption assay may be a more sensitive probe for the biological availability of the O1-antigen. This difference could be due to inconsistency in the LPS purification process that is absent in the phage-based assay, or due to a tight kinetic window where the proportion of O1-decorated LPS is changing rapidly around the two-hour timepoint.

The bile-dependent O1-depletion observed early upon exposure to these culture conditions indicated that cell growth rate alone may not be the determinant of O1-decorated LPS production because both bile-supplemented cultures produced less O1-decorated LPS prior to the anaerobic bile culture transitioning to a slower-growing state. Together, these results implicated a dynamic cellular response to bile early in the growth of the culture and anaerobic bile in subsequent overnight culture that negatively impacted *V. cholerae* O1-antigen production. The combination of bile and anaerobicity in overnight culture maintains low levels of O1-antigen production, and O1-antigen reduction negatively impacts ICP1 adsorption in all instances.

### RNA sequencing of *V. cholerae* grown in anaerobic, bile, and anaerobic bile conditions reveals transcriptional alterations in global cellular processes and pathogenesis genes

In order to identify transcriptomic changes underlying the observed reduction in O1-decorated LPS, we conducted RNA sequencing for all four culture conditions at both two-hour and overnight timepoints. We examined all pairwise comparisons of conditions where the O1-antigen was depleted in the experimental condition but not in the control condition (two-hour: aerobic bile vs aerobic and anaerobic LB, and anaerobic bile vs aerobic and anaerobic LB; overnight: anaerobic bile vs. all other conditions), reasoning that the overlap of the differentially expressed genes between conditions may identify candidates involved in the observed loss of the O1- antigen. Only eight genes were consistently differentially expressed across all seven pairwise comparisons from both timepoints (see Appendix). In all conditions comparing a bile-supplemented culture to a non-supplemented LB culture, we observed upregulation of genes involved in bile RND efflux (<u>r</u>esistance-<u>n</u>odulation-<u>d</u>ivision efflux superfamily) (*vexABCD*^33^), indicating that our transcriptomics experiment successfully recapitulated a known transcriptional response to bile in *V. cholerae*^34^ (Supplementary Figures S2D, S3D-S3E). However, none of the differentially expressed genes shared across all comparisons had direct logical connections to O1-antigen biosynthesis, indicating that instead of a simple conserved transcriptional mechanism, multiple factors may be involved in responses to both bile and anaerobicity that collectively impact O1-production.

Given the lack of clear candidate genes responsible for decreased O1-antigen levels, we used gene ontology to predict what metabolic processes were represented in the differentially expressed genes from each comparison. Gene ontology analysis revealed that genes encoding products involved in transmembrane transport dominated the pool of differentially expressed genes in O1-deplete conditions, both up- and downregulated (Figure 2A, Supplementary Figures S2, S3). More noticeably at the overnight timepoint, upregulated genes in O1-deplete conditions tended to be enriched in functions related to protein synthesis, e.g., translation, ribosome biogenesis, amino acid metabolism, and tRNAs, while the downregulated set was slightly enriched for genes involved in the regulation of gene expression and other/unclassified genes (Figure 2A, Supplementary Figures S2, S3). The transcriptomic data are consistent with a cell state where there may be adequate resources for transcriptional upregulation but limited resources to facilitate protein synthesis. Together, these data indicate that our transcriptomic profiling captured both previously-described and novel transcriptional adaptations to our tested conditions (see Appendix).

**Figure 2.**
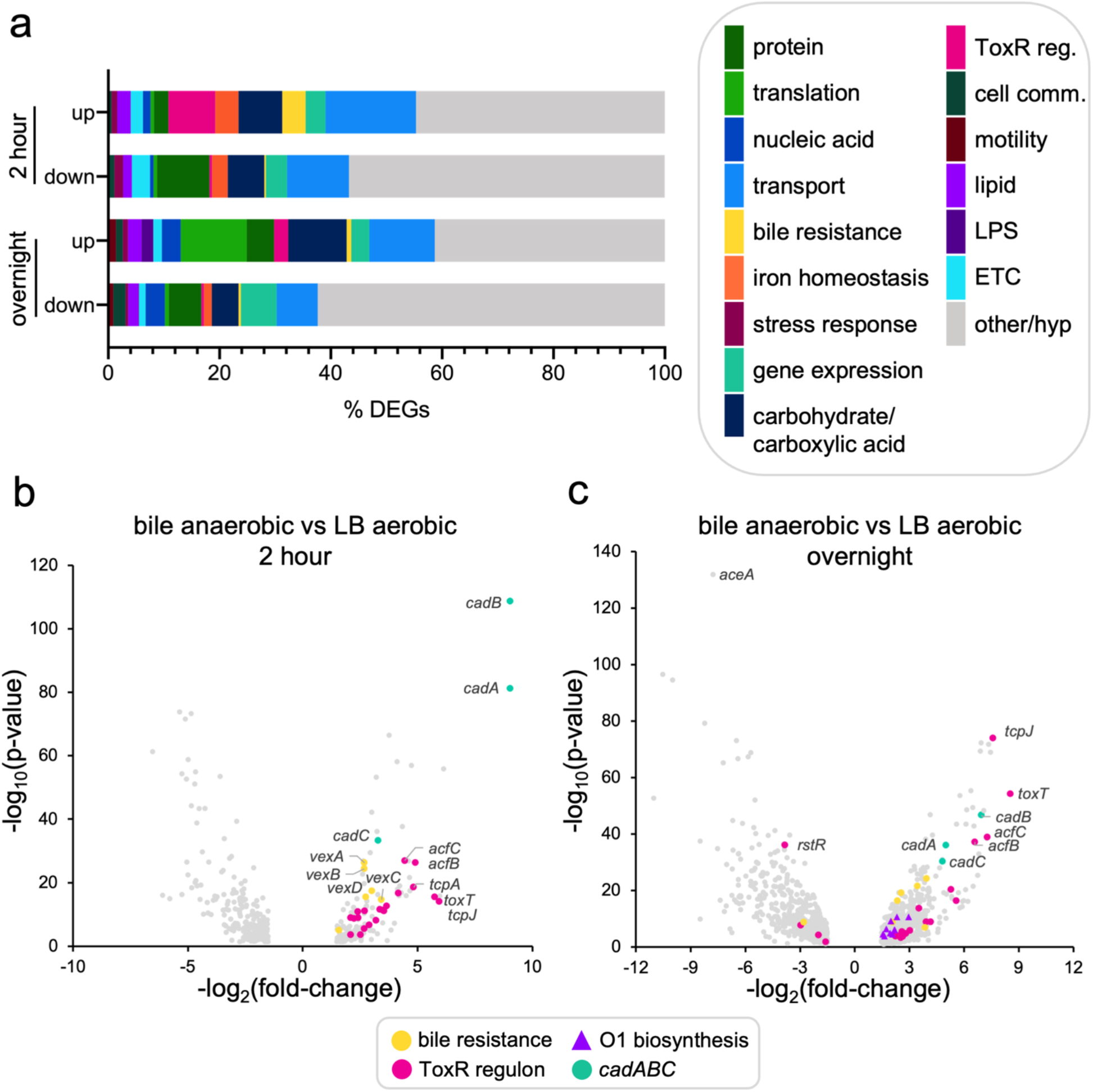
RNA sequencing of *V. cholerae* grown in anaerobic, bile, and anaerobic bile conditions reveals transcriptional alterations in global cellular processes and pathogenesis genes (**a**) Stacked bar graph representing the proportion of differentially expressed genes (DEGs) representing each considered gene ontology category. Differentially expressed genes from each individual comparison are pooled in this analysis. Categories represent gene ontology metabolic processes unless otherwise denoted. ‘ToxR reg.’: ToxR regulon, ‘cell comm.’: cell communication, ‘hyp’: hypothetical. (**b**) Two-hour and (**c**) overnight volcano plots of differentially expressed genes in anaerobic bile compared to aerobic LB conditions. Adjusted p-value p ≤ 0.05 and fold-change ≥ +/-1.5 was considered significant. Select gene categories of interest are highlighted in color and indicated in the legend. See Supplementary Figures S2 and S3 for volcano plots with designated gene ontology categories.

Many upregulated genes with the highest fold-changes in all pairwise comparisons were components of the ToxR regulon, specifically *toxT* (the transcriptional activator of downstream virulence cassettes CTX and TCP)^35^, the majority of TCP components^36^, and accessory colonization factors (*acfA*^37^*B*^38^*C*^39^) (Figure 2B-2C, Supplementary Figure S4). We observed activation of *ctxAB*^40^ and concordant downregulation of *rstR*^41^, the negative regulator of CTX, at the two-hour timepoint (Supplementary Figure S4B). Also indicative of ToxR regulon activity and bile resistance, *ompU* expression was increased in overnight comparisons, although the corresponding downregulation of *ompT*^30^ was only observed in one of the overnight comparisons (Supplementary Figure S4B). Bile and anaerobicity are both known activators of the ToxR regulon^42, 43^, which is consistent with the observation of its transcriptional upregulation in these conditions. ToxR induces a regulatory cascade for *V. cholerae* virulence gene expression induced in the human gut in the context of infection^12^, and has been shown to be highly transcriptionally upregulated in both rabbit and mouse animal models of cholera^44^. Interestingly, several other features of the distinct transcriptional profiles of *V. cholerae* in rabbit and mouse infections are also represented in the two-hour bile and overnight anaerobic bile culture conditions. This includes activation of a colonization factor (*vc1773*)^44^ encoded in *Vibrio* pathogenicity island 2 (VPI-2) and genes involved in the utilization of host-derived long-chain fatty acids found in cecal fluid (e.g., long chain fatty acid transport, acetyl CoA dehydrogenases, and glycerol transport and metabolism genes, which enable utilization of fatty acids)^44^ (Supplementary Figures S2-S3, Supplementary Data Sheet 3). At the overnight timepoint, we also observed upregulation of *almEFG* lipid A modification genes that are thought to be important in providing protection against antimicrobial peptides found in the gut^45, 46^ (Supplementary Figure S5). Taken together, the data sets suggest that *V. cholerae* producing less O1-decorated LPS in these culture conditions have also made global shifts in gene expression to a state remarkably similar to what has been observed in animal models of *V. cholerae* infection.

### Bile and anaerobicity alter the transcription of a subset of genes in the O1-biosynthetic cluster

We examined the O1-biosynthetic cluster itself to identify connections between growth conditions and transcription of the enzymes necessary to produce O1-antigen. To our surprise, seven consecutive genes within the O1-biosynthetic cluster (*gmd-wbeL*^47^) were significantly upregulated when comparing overnight anaerobic bile conditions to either aerobic or anaerobic LB (Figure 3A). This locus contains genes required for biosynthesis of both O1-antigen precursor components, perosamine and tetronate, as well as the required transport genes for O1- antigen assembly^47^. We were not expecting to observe a corollary transcriptional upregulation in this genomic region in conditions where less O1-decorated LPS was produced, so we proceeded to investigate if the observed transcriptional upregulation resulted in increased protein production. A representative gene from within (*wbeE*^47^) and outside (*wbeU*^47^) the upregulated set of genes was tagged with a C-terminal FLAG tag, and protein production was monitored by Western blot after overnight culture. While WbeU production remained similar across all culture conditions (consistent with no significant changes in expression detected by RNA-seq), there was a striking reduction in the amount of WbeE detected in cultures containing bile (Figure 3B- 3D). This suggested that the transcriptional upregulation within the O1-biosynthetic cluster was not indicative of enhanced protein production, but instead possibly indicating that cells were not producing enough of these enzymes and attempting to upregulate transcription to bolster low protein levels. Importantly, low abundance of WbeE was not sufficient to reduce O1-decorated LPS production, as similar levels of WbeE were produced in anaerobic and aerobic bile conditions and we observed complete O1 production in aerobic bile overnight culture. Taken with the RNA-seq data suggesting a global upregulation of genes encoding ribosomal and other translational components (Figure 2A, Supplementary Figures S2-S23), these results suggest that cells in anaerobic bile conditions may be adjusting to conditions with fewer resources for protein production by enhancing transcription as if to prepare for the introduction of more protein synthesis resources. These global changes alone could result in less O1-decorated LPS production via a reduction in the pool of enzymes essential for O1-biosynthesis. This would represent a mechanism of phage resistance tied to alterations in cellular processes that result in the restriction of ICP1 phage predation via the reduced presentation of the phage receptor.

**Figure 3.**
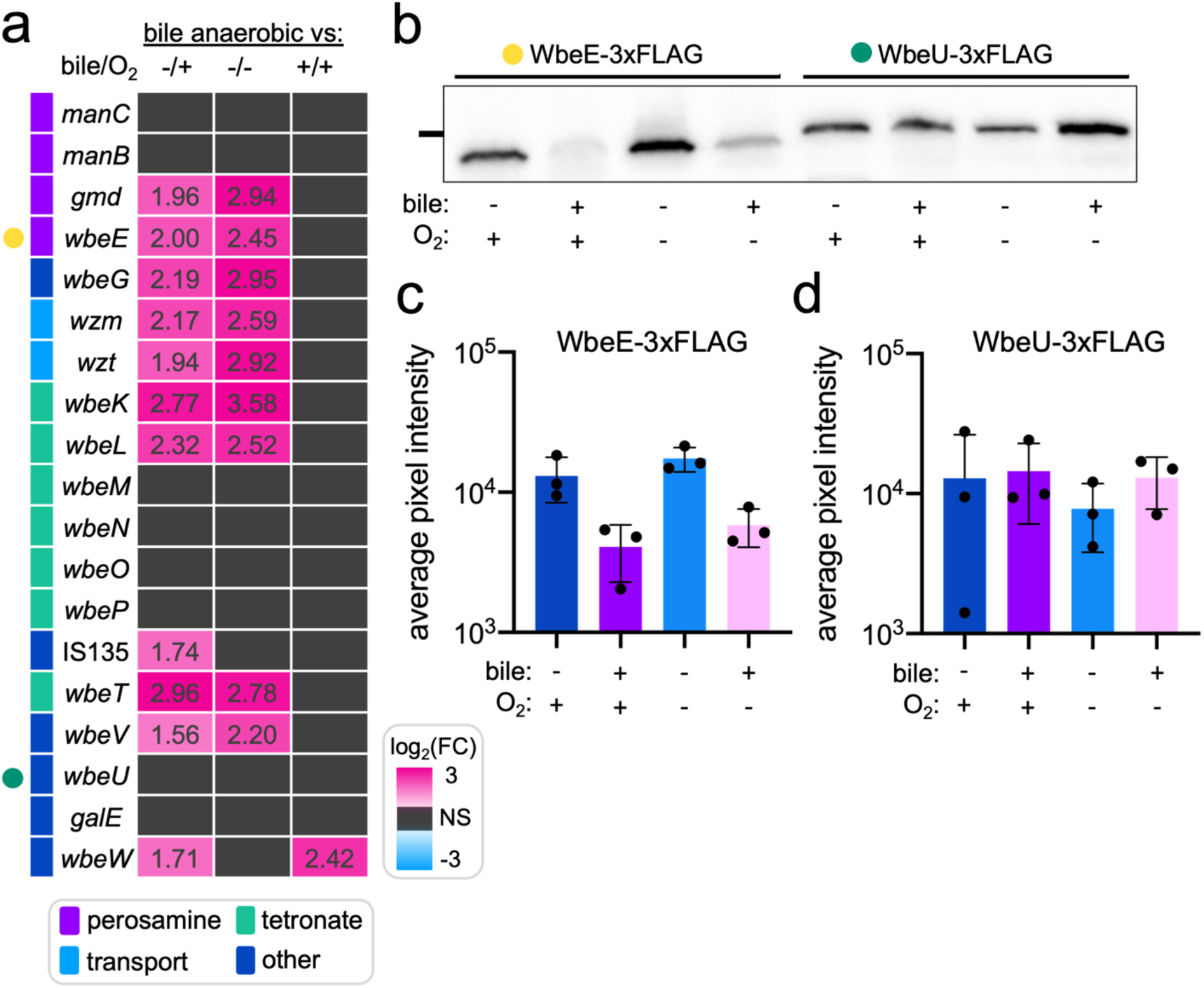
Bile and anaerobicity alter transcription of a subset of genes in the O1-biosynthetic cluster (**a**) Heatmap of log_2_(fold-change) for genes in the O1-biosynthetic cluster in overnight culture comparisons. log_2_(fold-change) values are listed in individual cells. Right legend gives colorimetric approximation of log_2_(fold- change), NS (black) denotes nonsignificant changes in transcript levels (p_adj_ > 0.05). Bottom legend provides function category for each gene (colored bars left of gene names). (**b**) Representative Western blot against FLAG- tag monitoring WbeE-3xFLAG and WbeU-3xFLAG protein expression in all combinations of culture conditions after overnight incubation. Left line represents 37kDa marker. (**c**) WbeE and (**d**) WbeU pixel intensity quantification for replicate Western blot experiments.

### The *cadABC* weak acid tolerance system is necessary for O1 biosynthesis in anaerobic bile conditions

From the pool of differentially expressed genes identified by RNA-seq, we selected and screened several candidate genes and systems with potential relevance to O1-antigen depletion in bile anaerobic conditions. We focused on genes implicated in bile resistance and bile-mediated transcriptional response (*vexABR*^48^, *leuO*^49^*, toxR*^50^) and systems with predicted utility in intestinal colonization (PTS carbohydrate transport system^51^, RTX toxin gene cluster^52^, *almEFG* lipid modification system^45^, citrate utilization, *cadABC* weak acid tolerance system^53, 54^), disabling a system’s regulator whenever possible. Surprisingly, these systems were all dispensable for growth in anaerobic bile conditions and nearly all mutants were able to synthesize as much O1-decorated LPS as wild-type *V. cholerae* (Supplementary Figure S10). A single exception was the Δ*cadC* mutant lacking the CadC sensor/transcriptional activator protein of a lysine-dependent weak acid tolerance system (Figure 4A). The Δ*cadC* strain produced LPS with even less O1-antigen decoration than wild-type *V. cholerae* in anaerobic bile conditions, and concordantly demonstrated a further reduction in ICP1 adsorption (Figure 4B, 4C, Supplementary Figure S6A). In response to low pH and lysine, CadC activates the expression of *cadA*, (a lysine decarboxylase that consumes protons and converts lysine to cadaverine) and *cadB* (a lysine/cadaverine antiporter)^53, 54^ (Figure 4A). RNA-seq data also implicated the importance of this gene cluster, as *cadB* and *cadA* were among the most significantly differentially expressed genes in both the two-hour and overnight timepoint comparisons (Figure 2B-2C, Supplementary Figure S3). To confirm that the activity of the *cadABC* system was required for maximum O1-antigen biosynthesis, we cultured a Δ*cadA* strain in anaerobic bile conditions and observed a decrease in O1-decorated LPS production on par with the Δ*cadC* strain (Supplementary Figure S7C, S7D). A further reduction in O1-antigen with loss of the weak acid tolerance system activity indicated that the *cadABC* system is functionally necessary for O1-decorated LPS production in anaerobic bile conditions.

**Figure 4.**
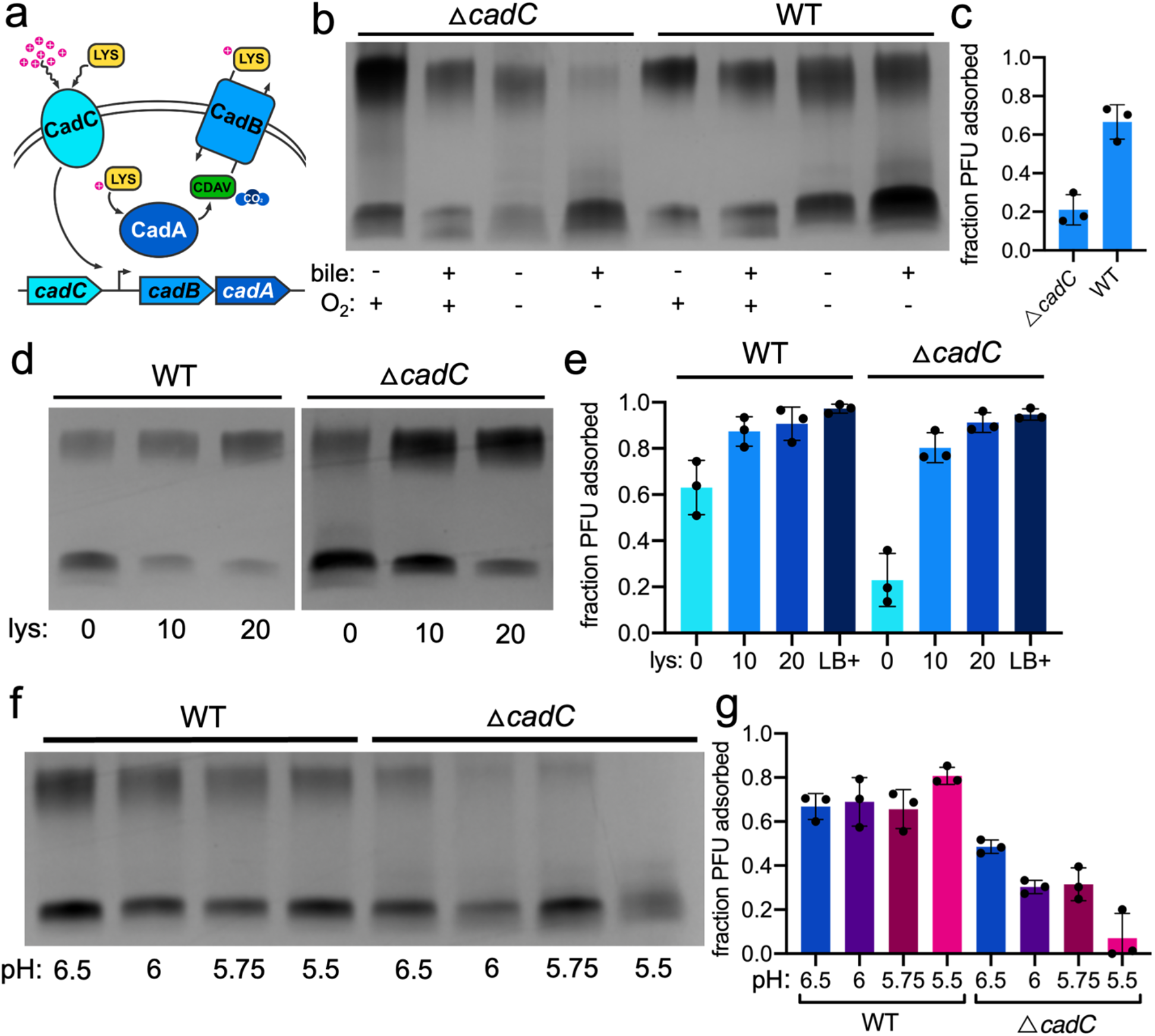
The *cadABC* weak acid tolerance system is necessary for O1 biosynthesis in anaerobic bile conditions (**a**) Schematic of the *cadABC* weak acid tolerance system. At neutral pH, membrane-bound CadC represses expression of *cadAB* (excluded from schematic). When CadC senses low extracellular pH and lysine, it activates expression of *cadB* (a lysine/cadaverine antiporter) and *cadA* (a lysine decarboxylase that consumes protons (magenta circles with (+)) to convert lysine (LYS, yellow) to cadaverine (CDAV, green) and carbon dioxide). CadABC activity neutralizes intra- and extracellular pH. (**b**) Purified LPS silver stain of wild-type (WT) and !ι*cadC V. cholerae* in all combinations of culture conditions. (**c**) Fraction of ICP1 adsorbed to *V. cholerae* grown overnight in anaerobic bile culture conditions. (**d**) Purified LPS silver stain of *V. cholerae* grown overnight in anaerobic bile culture conditions supplemented with L-lysine (lys, concentration denoted in mM). (**e**) Fraction of ICP1 adsorbed to *V. cholerae* grown overnight in anaerobic bile culture conditions supplemented with L-lysine (lys, mM) or aerobic LB (LB+). (**f**) Purified LPS silver stain of wild-type (WT) and !ι*cadC V. cholerae* grown overnight in anaerobic bile culture conditions, pH reduced with malic acid. (**g**) Fraction of ICP1 adsorbed to *V. cholerae* grown overnight in anaerobic bile culture conditions, pH reduced with malic acid.

While the loss of weak acid tolerance further depleted the production of O1-decorated LPS, none of the screened mutants were able to produce more O1-decorated LPS than wild-type *V. cholerae* in anaerobic bile conditions (Supplementary Figure S10). We tested chromosomal overexpression of *cadC* by synthetic induction in anaerobic bile conditions and did not observe any increase in O1-decorated LPS (Supplementary Figures S6B, S7A, and S7B). This could be indicative of limitations in cellular resources necessary for O1-antigen biosynthesis that are not replenished by the de-acidification activity of *cadABC*. Given that cells may lack robust protein synthesis capacity, we do not have direct evidence that synthetic overexpression can increase CadC protein concentration in these conditions. Taken together, these results suggest that *cadABC* activity is necessary to produce O1-decorated LPS in anaerobic bile conditions, but the reduction of O1-decorated LPS in wild-type cells may not be due to a simple lack of adequate CadABC system activity and could instead be a result of other metabolic changes interfering with cellular resources necessary for weak acid tolerance and O1 biosynthesis. Therefore, weak acid tolerance is also necessary for ICP1 infection in anaerobic bile conditions, as its activity is required for the production of the phage receptor.

### pH determines the degree of O1-decorated LPS production

Because synthetic overexpression of *cadC* did not result in the recovery of O1-decorated LPS, we sought to alternatively activate the *cadABC* system with the addition of exogenous lysine, a trigger of CadC activation. Initially, we were excited to observe that the addition of lysine appeared to have a positive, dose-dependent effect on O1-decorated LPS production in wild-type *V. cholerae*, with 20mM lysine supplementation of anaerobic bile *V. cholerae* cultures resulting in LPS comparable to what is observed in aerobic LB conditions (Figure 4D, 4E, Supplementary Figure S6G). To determine if the recovery of O1-decorated LPS was due to the activation of *cadABC*, we grew the Δ*cadC* strain with supplemental lysine, anticipating that lysine would no longer impact the production of O1-decorated LPS. Unexpectedly however, we observed the recovery of O1-decorated LPS with lysine supplementation for the Δ*cadC* mutant comparable to wild-type *V. cholerae* (Figure 4D, 4E, Supplementary Figure S6G). The Δ*cadA* strain also showed recovered production of O1-decorated LPS in anaerobic bile following supplementation with lysine (Supplementary Figure S7C, S7D, S6G). While the addition of lysine restored the production of O1-decorated LPS and correspondingly restored sensitivity to ICP1 adsorption (Figure 4E), bacterial growth did not increase (Supplementary Figure S9E-S9G). This indicates that general growth restriction is likely not the cause of O1-decoration loss and reduced phage adsorption in anaerobic bile conditions. Overall, these results suggested that the recovery of O1- antigen was not directly due to lysine-mediated activation of *cadABC*.

Lysine is a basic amino acid and *cadABC* is responsive to acidic conditions, so we considered the possibility that the addition of lysine was increasing the media pH, and this alkalization was the underlying cause of O1-antigen recovery during lysine supplementation. We first measured the pH of fresh and spent media from each relevant condition and confirmed that lysine supplementation increases pH from 6.8 to ∼8.5 in anaerobic bile culture (Supplementary Figure S7E). We therefore supplemented anaerobic bile cultures with arginine (another basic amino acid) at the same concentrations or adjusted the pH of the culture media with sodium hydroxide to match 10mM (∼pH 7.5) and 20mM (∼pH 8.5) basic amino acid supplementations and assessed the resulting LPS. Both arginine and sodium hydroxide supported robust recovery of O1-decorated LPS in both wild-type and Δ*cadC V. cholerae* (Supplementary Figure S7F, S7G), indicating that increased pH allows for complete O1-antigen production.

The pH of anaerobic bile media (both fresh and spent) was very close to neutral (∼pH 6.8) (Supplementary Figure S7E), while in the context of a human infection, *V. cholerae* experiences dramatic pH ranges in bile (∼pH 6-8)^55^ and stomach acid (∼pH 2-6)^56^. Because of the apparent essentiality of *cadABC* weak acid tolerance activity for O1-biosynthesis in anaerobic bile conditions, we hypothesized that reducing the pH in this condition could necessitate further increased CadAB activity for O1-production, meaning wild-type *V. cholerae* with active CadAB may continue to synthesize O1 until the pH is too low to be tolerated, while mutants in the weak acid tolerance system would lose O1-biosynthetic capacity at sufficiently low pH. We therefore examined the effects of low pH anaerobic bile conditions on LPS production in both wild-type *V. cholerae* and the Δ*cadC* strain. The wild-type strain was able to grow in bile anaerobic conditions as acidic as pH 5.5 (Supplementary Figure S9H) and did not exhibit any significant changes in O1-decorated LPS production at low pH (Figure 4F, 4G). While the Δ*cadC* strain grew at relatively similar rates as the wild-type at low pH (Supplementary Figure S9H), the LPS produced by this strain exhibited a further reduction in O1-antigen substitution with decreasing pH, where almost no detectable O1-antigen was produced at pH 5.5 (Figure 4F, 4G). We used an organic acid (malic acid) to reduce the pH of the media, however *cadABC* has been previously described as important for tolerance to both organic and inorganic acids in *V. cholerae*^57^.

Accordingly, when we reduced the pH of the media with an inorganic acid (hydrochloric acid) we observed the same dose-dependent reduction in O1-antigen in the absence of *cadABC* activity (Supplementary Figure S8A, S8B, S9I), suggesting that the phenotype is truly pH-dependent and independent of the origin of protons causing the acidity.

Because the acidity of the anaerobic bile condition appeared to be the determining factor in reducing O1-antigen production, we sought to determine if acidity was sufficient to reduce O1- decoration on LPS, or if bile supplementation and/or anaerobicity were also required. We measured growth and LPS production at low pH in anaerobic LB (Supplementary Figure S9J, S8C, S8D) and aerobic bile conditions (Supplementary Figure S9K, S8E, S8F) and did not observe any reduction in the degree of O1-decorated LPS produced by either wild-type or Δ*cadC* strains. These results suggest that bile, anaerobicity, and low pH combined reduce the production of O1-decorated LPS. Acid tolerance mediated by *cadABC* allows for partial O1-antigen production in these conditions, but the loss is significant enough to dramatically hinder the adsorption of ICP1 phage, providing an environmentally mediated mechanism of adaptive phage resistance.

## Discussion

To establish a thorough understanding of phage defense as it has evolved in native contexts, the interplay of many systems must be considered as a holistic picture. As the discovery and appreciation of novel phage defense systems rapidly accelerates, a more underappreciated route of inquiry dissecting the intertwined nature of phage defense and bacterial physiology has also recently gained appreciation Reversible environmental adaptation conferring phage resistance was recently described in an *E. coli* mouse model, where differential expression of biofilm genes and O-antigen ligase in susceptible bacterial hosts limited phage predation in the intestine^58^. *Streptomyces* species have been shown to convert to a cell-wall-deficient state when exposed to environmental pressures, including phage infection, and *E. coli* and *Bacillus subtilis* cultured in an osmoprotective environment also convert to a cell-wall-deficient state to resist phage infection by shedding the phage receptor^59^. The work presented here demonstrates a related phenomenon in *V. cholerae*, where exposure to signals relevant to the intestinal environment transiently alters the availability of O1-antigen decoration on LPS, negatively impacting the adsorption of a phage that requires the O1-antigen as a receptor. We investigated the cause of this reduced O1-antigen production and identified several routes by which the decrease may occur, including a reduction in O1-biosynthetic enzyme production and global changes in gene expression, indicating changes in the availability of protein translation and central carbon metabolism components required for constructing O1-antigen subunits. We also identified the essentiality of the weak acid tolerance system *cadABC* in the production of O1-antigen in conditions mimicking the intestinal environment (Figure 5).

**Figure 5.**
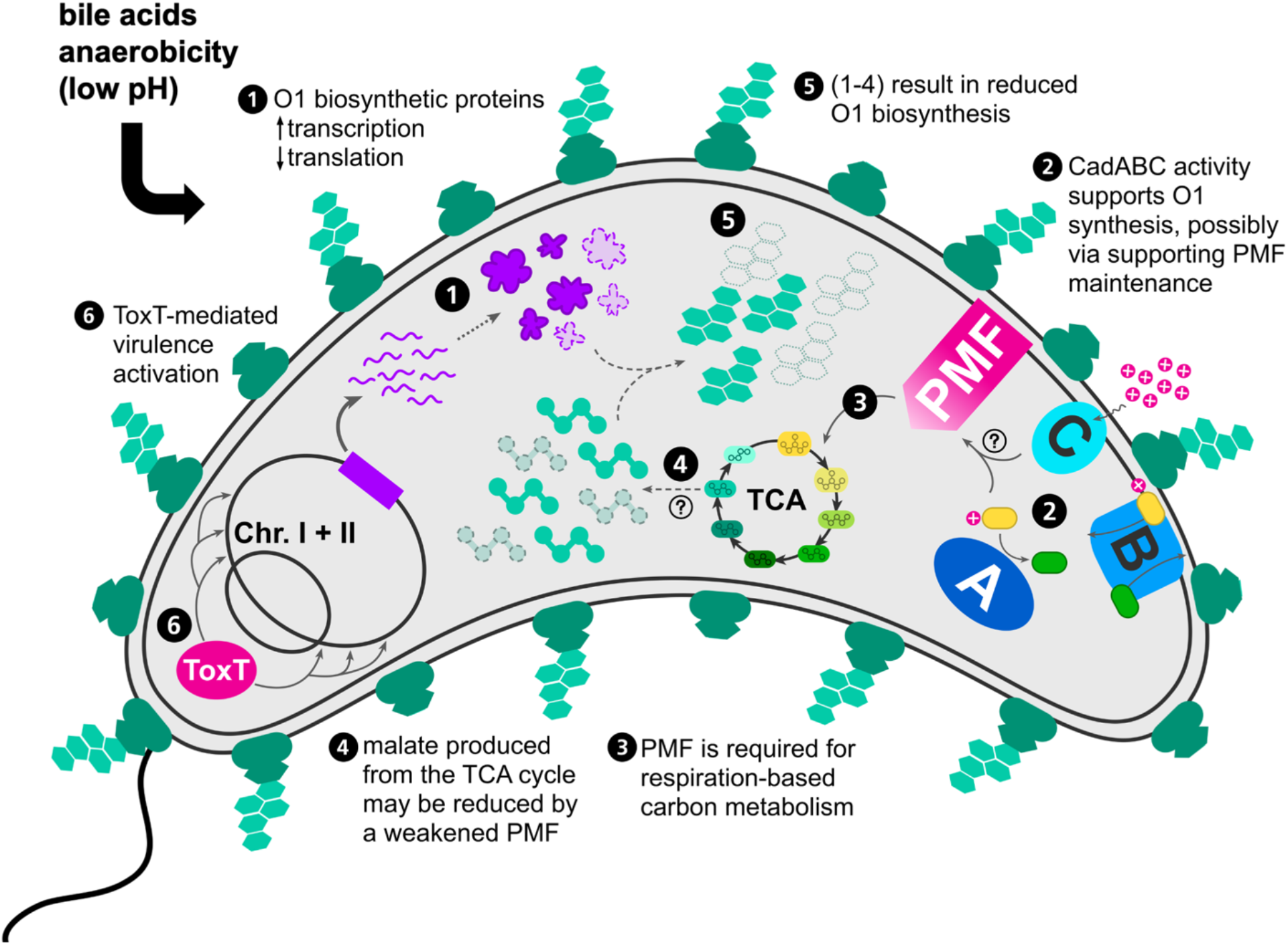
Mechanisms contributing to O1-antigen depletion in response to intestinal stimuli. (**1**) In O1-deplete conditions, pools of enzymes available for O1-antigen biosynthesis are reduced. This reduction is correlated with transcriptional upregulation of a putative operon within the O1-biosynthetic cluster (Figure 3). (**2**) The *cadABC* weak acid tolerance system is necessary for O1-antigen production in O1-deplete conditions (Figure 4). CadABC activity neutralizes cellular pH and may play a role in maintenance of proton motive force (PMF) in O1-deplete conditions, making its activity conditionally essential for robust carbon metabolism and O1-biosynthesis by proxy. (**3**) Cellular proton motive force (PMF) drives respiratory central carbon metabolism and ATP production. The tricarboxylic acid cycle (TCA) transfers electrons to cofactors (NAD+/FAD) that carry the electrons to the electron transport chain, which produces a proton gradient across the membrane (PMF) that is necessary to synthesize ATP from ADP via ATP synthase (utilizing either oxygen or alternative terminal electron acceptors). The recycling of cofactors from electron transport for continued utilization in the TCA cycle is essential for continued carbon metabolism. (**4**) Malate is a product of the TCA cycle and an essential precursor for O1-antigen biosynthesis. One potential result of weakened PMF is a reduction in pools of carbon metabolites such as malate. (**5**) Reduction in availability of O1-biosynthetic enzymes and putative reductions in malate pools result in a decreased capacity to synthesize O1-antigen. (**6**) In the conditions tested, O1-depletion occurs in the context of ToxT-mediated activation of a canonical *V. cholerae* pathogenesis transcriptional cascade.

This study is the first to describe altered transcription of O1-biosynthetic genes in response to intestine-associated stimuli coupled with the activation of the ToxR virulence transcriptional cascade. *In vivo* transcriptomic experiments in animal models did not detect differential expression of the O1-biosynthetic cluster^44^. Previous characterizations of *V. cholerae’s* response to bile acids and anaerobicity individually were focused on relatively limited microarray^34^ and 2D gel proteomic studies^60^, respectively. While our work is the first (to our knowledge) to apply RNA sequencing to assess the synergistic effects of bile and anaerobicity on the *V. cholerae* transcriptome, our observations align well with previous analyses of these environmental factors and allowed for rich contextualization of some subtle trends in our transcriptomic data (see Appendix). Reminiscent of what we observed in anaerobic bile conditions, previous transcriptional analyses of *V. cholerae* in mouse and rabbit intestinal models identified the upregulation of pathogenesis, transport, and regulatory genes^61^ coupled with transcriptional changes to fatty acid and glycerol metabolism, and activation of cryptic colonization factors in VPI-2 (*Vibrio* pathogenicity island-2), but no transcriptional changes in the O1-biosynthetic cluster^44^. One potential explanation for this discrepancy could be differences in environmental structure. Growing bacteria in liquid culture represents a physically homogeneous mixture, while the intestinal environment is spatially complex. Spatial complexity has been shown to play a role in the stable coexistence of phages and bacteria^62^ and is likely to contribute to the development of subpopulations with different transcriptional profiles. Such nuances would be lost in averaging transcriptome data for the entire population harvested from animal models. This could also help contextualize the surprising observation of O1-antigen reduction in anaerobic bile conditions considering the essentiality of the O1-antigen for intestinal colonization. It is possible that only some *V. cholerae* in an *in vivo* gut environment respond to collective bile, low pH, and anaerobic signals by reducing O1-antigen-decorated LPS, while other subpopulations experience different intensities of these signals and produce a different transcriptional response. This would allow for some subpopulations to express the optimal amount of O1-antigen for intestinal colonization, while other subpopulations potentially sacrifice some degree of colonization efficiency for protection from phage predation. Alternatively, O1-antigen expression may be important for the establishment of intestinal colonization but more dispensable later during infection, allowing for a population to efficiently colonize and then shift to a more phage-resistant state after initial establishment.

Previous work has also described transcriptional activation of *cadA* in rabbit and mouse cholera models^57^. Although there was no observed colonization defect in a *cadA* mutant strain, there was an observed colonization advantage for acid-adapted *V. cholerae*, in which the *cadABC* system plays a pivotal role. We observed a very limited amount of O1-antigen produced by the *cadA* and *cadC* mutants in anaerobic bile conditions, which would be expected to significantly reduce colonization efficiency. Because previous observations of *cadA* mutants *in vivo* did not indicate colonization deficiency, this further supports the hypothesis that the *in vivo* environment provides niches with a variety of conditions that allow for *V. cholerae* to thrive with different degrees of O1-antigen production. We also observed a pH-dependent depletion of O1-decorated LPS under both organic and inorganic acid stress in the absence of weak acid tolerance, consistent with the previously identified role of *cadA* in the *V. cholerae* acid tolerance response to both types of acid^57^. Overall, we identified that weak acid tolerance via *cadABC* is required for O1-biosynthesis under anaerobic bile conditions, suggesting that *cadABC* upregulation observed *in vivo* may be related to enhancing colonization efficiency by supporting O1-antigen biosynthesis. The requirement for *cadABC* could be due to the differential accumulation of metabolic byproducts in bile and the absence of oxygen that feed into O1-biosynthesis, or due to an undiscovered transcriptional connection between the regulation of weak acid tolerance and O1-biosynthesis.

While we identified several potential mechanisms by which O1-antigen production could be decreased in anaerobic bile conditions, we have yet to elucidate a direct link between O1- decoration and weak acid tolerance mediated by *cadABC*. In general, bacterial acid tolerance systems have been hypothesized to play essential roles in the maintenance of proton motive force (PMF) in extreme pH conditions^63^. In *V. cholerae*, the hydrochloric-acid-specific tolerance system (*clcA*) relies on tightly regulated expression of chloride channels that exchange intracellular chlorine for hydrogen ions to avoid toxic membrane hyperpolarization (termed an “electrical shunt”). These channels must be quickly silenced in more alkaline pH conditions to avoid PMF disruption, and may act in tandem with amino acid decarboxylation acid tolerance systems like *cadABC*^63, 64^. The impact of anaerobic bile conditions on PMF is unclear, but there is a hypothetical role for *cadABC* activity in the maintenance of PMF and a logical link to O1- biosynthesis. CadA decarboxylates lysine and consumes protons in the process, and CadB consumes protons while importing lysine. These activities can increase cytoplasmic and periplasmic pH. If there is more decarboxylation of cellular lysine, the cytoplasmic pH neutralizes more rapidly and produces a stronger PMF. PMF is required for energy production via ATP synthase in the last step of cellular respiration. Energy production from ATP synthase also recycles metabolic cofactors (e.g. NAD+/FAD), which allows for continued production of central carbon metabolism components and ATP. In anaerobic conditions, *V. cholerae* likely relies on anaerobic respiration with alternative electron acceptors for energy metabolism^65, 66^. A weakened PMF can slow down the entire carbon cycle due to reduced turnover of cofactors^67^. Components of central carbon metabolism serve as the building blocks for O1-antigen subunit biosynthesis (perosamine is synthesized from D-fructose 6-phosphate and tetronate is synthesized from malate^47^), so alterations in the PMF could negatively impact the availability of these components for O1-antigen biosynthesis. We hypothesize that the *V. cholerae cadA* and *cadC* mutants produce even less O1-antigen in anaerobic bile conditions because they are unable to strengthen the PMF with weak acid tolerance activity, resulting in a further hindrance of central carbon metabolism and reduction of O1-subunit biosynthesis components (Figure 5). Taken together, this work suggests a cryptic role for acid tolerance as a determinant of phage infection by proxy of supporting biosynthesis of the O1-antigen phage receptor.

Membrane surface modification in *V. cholerae* appears to be a common strategy to afford resistance to many different biological threats. *V. cholerae* responds to bile stimulus with a ToxR-regulon-mediated exchange of outer membrane porin OmpT for OmpU, a different porin with an important role in bile resistance^12^. *V. cholerae* lipid A is also modified via glycine addition, facilitating resistance to antimicrobial peptides during intestinal pathogenesis^45^. Exchange of OmpT and unmodified lipid A for OmpU and modified lipid A first occurs at the transcriptional level, but clearance of existing OmpT and unmodified lipid A in the outer membrane is achieved by the production of outer membrane vesicles (OMVs), which are extrusions of accumulated phospholipids, LPS, and proteins from the outer membrane^68^. This OMV-mediated surface exchange is also proposed to confer phage resistance: OMVs can act as decoys carrying phage receptors where phage may bind, interfering with receptor binding on viable *V. cholerae* cells. This effect has been demonstrated with vibriophages including ICP1, where exposure to high concentrations of OMVs modestly reduced phage titers^69^. With the caveat that we did not observe differential expression of genes involved in OMV production in our RNA-seq analysis, OMV-mediated surface exchange coupled with decreased O1-antigen biosynthesis (which is a likely consequence of the observed depletion of O1-biosynthetic proteins (Figure 3B-3D)) could result in the reduction of O1-decorated LPS we observed from *V. cholerae* cells in anaerobic bile culture. This would not only allow for the production of decoys but also provide a mechanism for shedding a substantial amount of phage receptor from the cell surface, hypothetically adding another mode of OMV-mediated phage resistance. However, it is currently unknown if OMV production plays a role in the reduction of O1-decorated LPS.

This work explores a physiological response to signals from *V. cholerae*’s natural environment that reduces O1-antigen decoration on LPS, an essential moiety for efficient intestinal colonization and for lytic phage predation. Studying the impact of these signals isolated from the *in vivo* environment allowed us to capture direct and relatively homogeneous responses to bile and anaerobicity that would not be possible in the heterogeneous *in vivo* intestinal environment. This response afforded protection against phages targeting the O1-antigen as a phage receptor, such as ICP1, without eliminating the antigen altogether. Reduction in O1-decorated LPS appeared to be a transient response to bile and anaerobicity, which are signals that are transiently encountered by *V. cholerae* while infecting a human host. As *V. cholerae* travels through the diverse “biomes” of the human digestive tract, the bacteria can temporarily alter their surface composition to respond to environmental changes without permanent disruption of genetic sequence. Such transient reduction in phage susceptibility could help explain why we observe both phages and phage-susceptible *V. cholerae* in cholera patient stool. The results underscore the importance of interrogating biological interactions in their environmental context.

Incorporating environmental signals into standard laboratory culture conditions is not equivalent to studying complex environmental frameworks, but it allows for the impact of such signals to be interrogated in detail. The environmental context appears to be particularly important for predatory biological interactions like phage infections, which drive the rapid evolution of both phage and bacteria in nature.

## Materials & Methods

### Strain construction

*V. cholerae* mutant strains with spectinomycin marker genes were constructed from E7946 (wild-type background) via natural transformation of linear PCR products^70^ that were synthesized by splicing by overlap extension PCR^71^ containing a spectinomycin resistance cassette or desired insertion sequence with 1kB homologous sequence flanking each side. Natural transformants were selected on LB supplemented with spectinomycin (100μg/mL) for knockouts or kanamycin (75μg/mL) for insertions in the *lacZ* locus, and modifications were verified by Sanger sequencing. *V. cholerae* marker-less mutants were constructed by the pCVD442-lac method as previously described^24^.

### Bacterial & phage growth conditions

*V. cholerae* E7946 and derivatives (a complete list of strains can be found in Supplementary Table 1) were cultured from 20% glycerol stocks stored at -80°C by streaking onto LB agar plates incubated at 37°C overnight. Plates were used to inoculate LB liquid media (Fisher Scientific), and liquid cultures were incubated at 37°C with aeration. Bile-supplemented cultures contained 0.5% purified dehydrated ox bile (Millipore-Sigma) in LB unless otherwise specified. Anaerobic media was degassed overnight and aliquoted into Hungate culture tubes (Chemglass Life Sciences CLS420801) in an anaerobic chamber (Coy Lab Products, 2.3% hydrogen, <30ppm oxygen). Growth assays were conducted by growing cultures to saturation in aerobic LB, then back-diluting to OD_600_=0.05 and allowing for growth in all media conditions for either 2 hours or overnight (∼16 hours). Anaerobic media was inoculated with a small volume of saturated aerobic culture to an OD_600_=0.05 (20-50μl per 5mL culture) diluted in 200μl of LB and injected through the rubber stoppers with a 23g syringe. For strains with chromosomal expression constructs, cultures were grown to saturation aerobically in LB supplemented with 1mM isopropyl β-D-1-thiogalactopyranoside (IPTG) and 1.5mM theophylline, then back-diluted as described above in media containing the same concentrations of IPTG and theophylline.

Cultures were supplemented with varying amounts of a 1M liquid stock of malic acid, L-lysine, L-arginine (Millipore-Sigma), hydrochloric acid, or sodium hydroxide (Fisher Scientific) to the designated concentration or final media pH.

Phage strains were propagated via soft agar overlay method on wild-type *V. cholerae* (for ICP1) and Δ*wbeL* (O1-) *V. cholerae* (for ICP2) (0.5% agar in LB, grown at 37°C for 6 hours). Phages were harvested in STE buffer (1M NaCl, 200mM Tris-HCl, 100mM EDTA) overlayed on confluent lysis plaque plates, rocking overnight at 4°C. Buffer was then collected, chloroform-treated and cleared by centrifugation at 5,000 x g for 10 minutes. Phage in the cleared lysate were then concentrated to a high titer by polyethylene glycol precipitation as described^72^.

### LPS purification

Bacteria were grown as described in “Bacterial & phage growth conditions” and pellets were harvested via centrifugation at 5,000 x g for 3 minutes. LPS was purified via hot aqueous phenol extraction method^73^: pellets were lysed in 200μl lysis buffer (2% 2-mercaptoethanol (BME), 10% glycerol in 0.1M tris-HCl pH6.8), boiled for 15 minutes, then treated with 25 units (5μl) RNase-A, 25 units (5μl) DNase-I, and 10 units (10μl) Proteinase K overnight at 37°C. Following overnight incubation and samples were incubated at 59°C for 4 hours. Next, an equal volume of cold tris-saturated phenol (pH 6.8) was added, and samples were mixed and incubated at 65°C for 15 minutes. Samples were then treated with 2.5 volumes cold diethyl ether, mixed thoroughly by inversion, then phase-separated by centrifugation at 20,000 x g for 10 minutes. The organic phase was extracted and the phenol extraction process was repeated a second time or until the organic phase was clear. Growth assays and purifications were performed in biological triplicate.

### LPS silver stain gels

Purified LPS was mixed with Lamelli sample buffer (Bio-Rad) to a 1x final concentration, boiled for 10 minutes, then run alongside a standard serial dilution of LPS purified from aerobically-grown wild-type *V. cholerae* on 4-12% Bis-Tris Criterion XT gels (Bio-Rad) at 165V for 40 minutes. Gels were washed with water then fixed overnight with rocking at 4°C in a gel fix solution (40% ethanol and 10% acetic acid). Gels were stained the following day using the sensitizing and staining solutions from the SilverQuest Silverstain kit (Invitrogen) followed by development with a modified developer (30% sodium carbonate and 0.05% formaldehyde) and stop solution (5% acetic acid). Gels were then imaged on an EZ Dock gel imager (Bio-Rad) with the white light sample tray. Image quantification was conducted in ImageJ. The intensity of the standard serial dilution of *V. cholerae* LPS (total lane intensity including both O1 and lipid A/core components) was used to generate a standard curve and calculate an intensity-standardized volume for each sample, then a second gel with those volumes of purified LPS were run and silver stained for final images and image analysis.

### Adsorption assays

*V. cholerae* cultures were grown as described in “Bacterial & phage growth conditions”. OD_600_ was measured and used to calculate a sample volume equivalent to OD_600_=0.3 in 1mL. Samples were pelleted by centrifugation at 5,000 x g for 3 minutes, spent media was aspirated and the pellet was washed with 1mL LB, re-pelleted by centrifugation, then resuspended in 1mL LB supplemented with 10mM magnesium chloride. Samples were heat-killed at 55°C for 10 minutes, cooled at room temperature for 5 minutes, then infected with phage at an MOI=0.01 (MOI=multiplicity of infection, calculated based on the number of viable cells in the sample volume pelleted prior to heat-killing). Infected samples were incubated at 37°C with aeration for 30 minutes to allow for complete adsorption. Next, samples were treated with 20μl of chloroform, vortexed, then separated by centrifugation at 5,000 x g for 15 minutes. Lysate was then serially diluted and used in plaque assays with a *V. cholerae* grown to an OD_600_=0.3. A parallel control was included with no cells (magnesium-supplemented LB heated in the heat-kill treatment prior to phage addition) to enumerate the exact phage input titer. Experiments were conducted in technical duplicate and biological triplicate.

### Growth curves

*V. cholerae* was grown to OD_600_∼1 in LB at 37°C with aeration. The culture was then back-diluted to OD_600_=0.05 in aerobic or anaerobic media, with or without 0.5% bile acid supplementation. Anaerobic cultures were quickly aliquoted into a clear 96-well plate and then sealed thoroughly with an optical seal (Bio-Rad), Aerobic cultures were similarly added to the other half of the plate, which remained unsealed. OD_600_ was measured over time in the 96-well plate shaking at 37°C in a plate reader (SpectraMax i3x). For paired CFU samples, separate cultures in aerobic and anaerobic tubes were started in parallel and harvested at the designated timepoints throughout the experiment. Growth curves were performed in technical and biological triplicate, a representative technical triplicate average is displayed in Figure 1I and biological replicates in Supplementary Figure S1H.

### Recovery assay

*V. cholerae* were grown overnight in anaerobic bile conditions as described in “Bacterial & phage growth conditions”. Cultures were harvested by centrifugation at 5,000 x g for 3 minutes, spent media was aspirated, and cell pellets were resuspended in an equal volume of fresh aerobic LB media. Cultures were then incubated at 37°C with aeration, and samples were taken at the designated timepoints to harvest LPS or conduct adsorption assays. Recovery assays were performed in technical duplicate and biological triplicate.

### RNA sequencing and analysis

#### RNA isolation

Cultures were grown as described in “Bacterial & phage growth conditions” section and cell pellets (approximately 8 x 10^7^ CFU) were harvested by centrifugation at 5,000 x g for 3 minutes. Pellets were resuspended in 200μl TRI Reagent (Millipore-Sigma) and incubated for 5 minutes at room temperature before the addition of 40μl chloroform. Samples were mixed thoroughly, incubated for 10 minutes at room temperature, then phase-separated by centrifugation at 12,000 x g for 15 minutes at 4°C. The aqueous phase was extracted and mixed thoroughly with 110μl isopropanol and 11μl 3M sodium acetate (pH 7.4), then incubated at room temperature for 10 minutes. Samples were phase-separated again by centrifugation as described above, then the aqueous phase was extracted, washed with 1mL 75% ethanol, then centrifuged again to pellet RNA. Ethanol was aspirated, and the residual was evaporated at 65°C prior to pellet resuspension in 40μl DEPC treated water (Growcells.com).

#### RNA sequencing

Approximately 1 µg of RNA per sample was submitted to the Microbial Genome Sequencing Center for 2x150 paired-end sequencing and RNA sequencing was carried out according to the Microbial Genome Sequencing Center protocols. Briefly, samples were DNAse treated with Invitrogen DNAse-I (RNAse free) and library preparation was performed using Illumina’s Stranded Total RNA Prep Ligation with Ribo-Zero Plus kit and 10bp IDT for Illumina indices. Sequencing was done on a NovaSeq 6000 giving 2x51bp reads.

Demultiplexing, quality control, and adapter trimming was performed with bcl-convert (v4.0.3).

#### RNA sequence analysis

Demultiplexed and trimmed fastq files were mapped to the *V. cholerae* E7946 reference genomes (Accessions: NZ_CP024162 & NZ_CP024163) using bowtie2 (v 2.3.5.1) and the resulting .bam files were sorted using samtools (v 1.9). Sorted .bam files were assembled into transcripts with Stringtie (v 2.1.7) with options (-eB) to output in ballgown format. Ballgown files for all samples were converted into hypothetical read counts for each transcript using the Stringtie-associated prepDE.py python script. The resulting transcript count matrix was imported into DESeq2 (v 1.34.0) for further analysis in R.

To perform QC analysis and to determine what appropriate comparisons could be generated from the dataset, the transcript count matrix was transformed according to the regularized log transformation (rlog) in DESeq2 and principal component analysis (PCA) was performed using PCAtools (v2.6). Dendrograms with heatmaps of the rlog transformed transcript count matrix were constructed using the R package pheatmap (v 1.0.12).

Differential expression analysis was performed with DESeq2, using the contrast function to generate pairwise comparisons of samples within a single timepoint according to their media conditions and aerobicity, to allow for a single-factor DESeq design formula. Genes were considered to be differentially regulated with a log_2_ fold-change of ± 1.5 and above a minimum adjusted p-value of 0.05. Volcano plots were generated from DESeq values in Microsoft Excel (Supplementary Data Sheet 1) and differentially expressed genes were colored according to prediction function from a combination of GO gene ontology prediction^74, 75^ and manual curation.

### Western Blot

*V. cholerae* cultures were grown overnight aerobically and anaerobically in LB with and without 0.5% bile acid supplementation. OD_600_-standardized pellets were harvested by centrifugation at 5,000 x g for 3 minutes, washed once with cold PBS, then resuspended on ice in cold lysis buffer (50 mM Tris, 150 mM NaCl, 1 mM EDTA, 0.5% Triton X-100, 1x Pierce™ Protease Inhibitor Mini Tablet (Thermo)). Protein concentration was determined using a Pierce BCA Protein Assay Kit (Thermo) and 30μg total protein per sample was mixed with Laemmli sample buffer (10% 2- mercaptoethanol added) to a final concentration of 1x. Samples were boiled at 99°C for 10 minutes, run on Any-kD TGX-SDS-PAGE gels (Bio-Rad) at 200V for 25 minutes, then transferred onto nitrocellulose membranes using a Transblot Turbo Transfer system (Bio-Rad). Following overnight blocking in 5% milk, membranes were incubated with a primary rabbit anti-FLAG antibody (Invitrogen) diluted 1:1500 for 3 hours. Signal detection was conducted with a goat anti-rabbit-HRP secondary antibody followed by development with Clarity ECL Substrate (Bio-Rad). Blots were imaged on Chemidoc XRS Imaging System (Bio-Rad). Band quantification was conducted using ImageJ. Western blots were conducted in biological triplicate.

### Data availability

Transcriptomic data from the RNA sequencing experiment are deposited in the Sequence Read Archive under the BioProject accession PRJNA954450.

## Supporting information

Supplementary Data Sheets 1-3

## Acknowledgements & Funding Sources

We would like to thank current and former members of the Seed Lab for useful discussions and feedback, particularly Zach Barth for thoughtful ideas and manuscript editing. Thank you to Arash Komeili and the Komeili Lab for access to their anaerobic chamber and for helpful feedback. We would also like to thank James Olzmann, Jeff Cox, and Kathleen Ryan for their guidance, suggestions, and support. This work was supported by a National Science Foundation Graduate Research Fellowship [2018257700 to D.T.D.] and by the National Institutes of Health [R01AI127652 to K.D.S, R01AI153303 to K.D.S]. Its contents are solely the responsibility of the authors and do not necessarily represent the official views of the National Institute of Allergy and Infectious Diseases or NIH. K.D.S. holds an Investigators in the Pathogenesis of Infectious Disease Award from the Burroughs Wellcome Fund.

## Notes

### Competing Interest Statement

The authors have declared no competing interest.

