## Supplementary material for "Adaptation to bile and anaerobicity limits *Vibrio cholerae* phage adsorption": Figs S1-S11 and Table S1

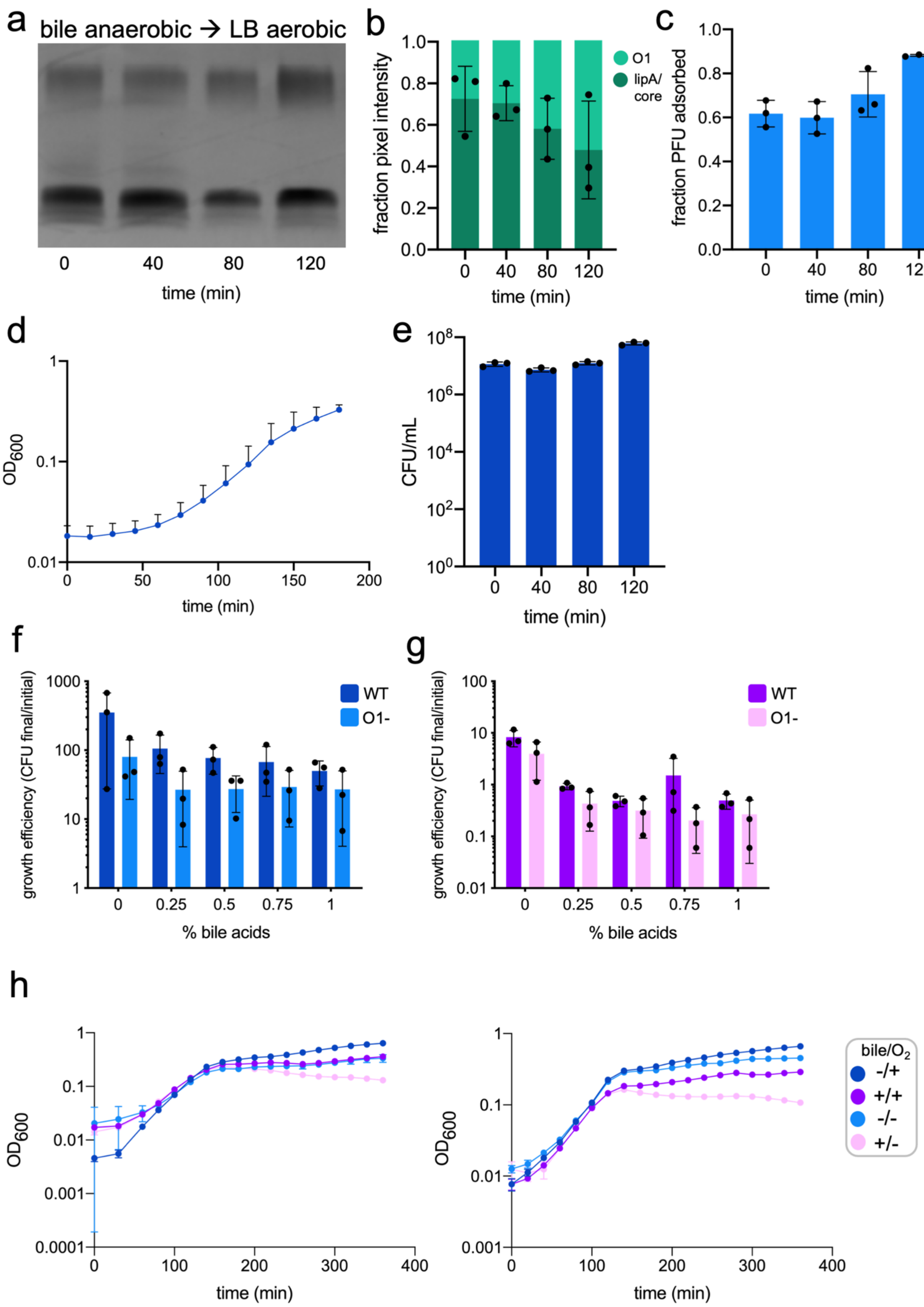

a
bile anaerobic → LB aerobic
0
40
80
120
time (min)
b
fraction pixel intensity
O1
lipA/ core
time (min)
c
fraction PFU adsorbed
time (min)
d
$OD_{600}$
time (min)
e
CFU/mL
time (min)
f
growth efficiency (CFU final/initial)
WT
O1-
% bile acids
g
growth efficiency (CFU final/initial)
WT
O1-
% bile acids
h
$OD_{600}$
time (min)
bile/$O_2$
-/+
+/+
-/-
+/-

**Supplementary Figure S1**

(**a**) Purified LPS silver stain of *V. cholerae* grown overnight in anaerobic bile conditions, then recovered in aerobic LB, time in minutes post-recovery. (**b**) Average fraction of pixel intensity contributed by O1 (light green) and lipid A/core (dark green) quantifying LPS purification and silver stain replicates represented in (a). (**c**) Fraction of ICP1 adsorbed to *V. cholerae* anaerobic bile overnight cultures at timepoints during recovery in aerobic LB. (**d**) $OD_{600}$ and (**e**) colony-forming units (CFU) of *V. cholerae* anaerobic bile overnight cultures recovering in aerobic LB. (**f**) Growth efficiency of wild-type (WT) and O1-antigen-null (O1-, Δ*wbeL*) *V. cholerae* grown aerobically in increasing concentrations of bile acids. Growth efficiency is calculated as $CFU_{final}/CFU_{initial}$. (**g**) same as (f) but in anaerobic conditions. (**h**) Biological replicates (each in technical triplicate) of growth monitored by $OD_{600}$ over time in all combinations of culture conditions represented in Figure 1I.

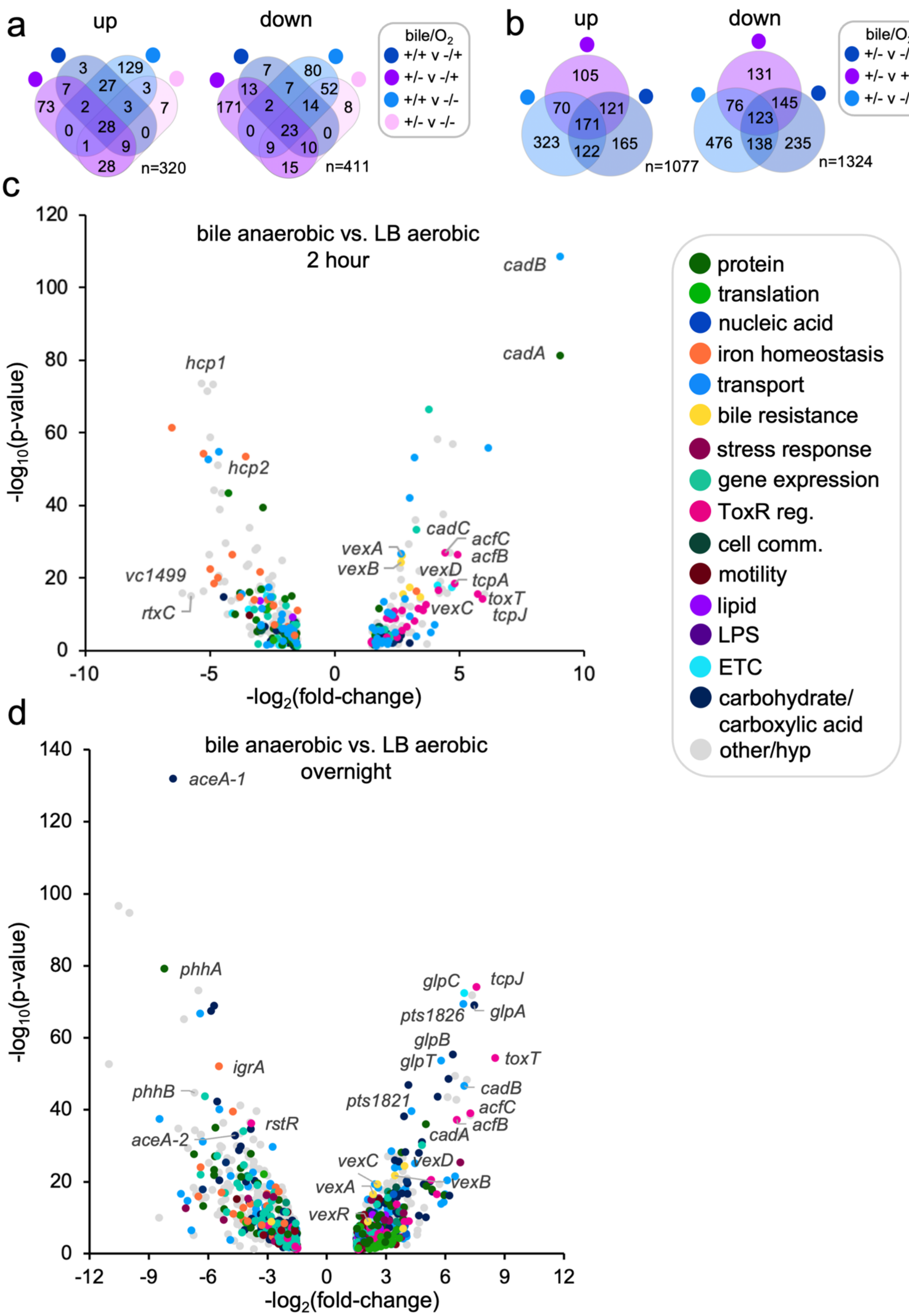

a
up
down
3
129
7
27
3
73
2
3
7
0
28
0
1
9
28
n=320
7
80
13
7
52
171
2
14
8
0
23
0
9
10
15
n=411
bile/O2
+/+ v -/+
+/- v -/+
+/+ v -/-
+/- v -/-
b
up
down
105
70
121
171
323
122
165
n=1077
131
76
145
123
476
138
235
n=1324
bile/O2
+/- v -/+
+/- v +/+
+/- v -/-
c
bile anaerobic vs. LB aerobic
2 hour
-log10(p-value)
-log2(fold-change)
cadB
cadA
hcp1
hcp2
cadC
acfC
acfB
vexA
vexB
vexD
tcpA
toxT
tcpJ
vexC
vc1499
rtxC
protein
translation
nucleic acid
iron homeostasis
transport
bile resistance
stress response
gene expression
ToxR reg.
cell comm.
motility
lipid
LPS
ETC
carbohydrate/
carboxylic acid
other/hyp
d
bile anaerobic vs. LB aerobic
overnight
-log10(p-value)
-log2(fold-change)
aceA-1
phhA
igrA
phhB
rstR
aceA-2
glpC
tcpJ
pts1826
glpA
glpB
glpT
toxT
cadB
pts1821
acfC
acfB
cadA
vexC
vexD
vexA
vexB
vexR

**Supplementary Figure S2**

(**a**) Venn diagram depicting differentially expressed genes shared between pairwise comparisons at two hours. "n" denotes total number of genes. (**b**) same as (a) at the overnight timepoint. (**c**) Two-hour and (**d**) overnight volcano plots of differentially expressed genes in anaerobic bile compared to aerobic LB conditions. Plots represent the same data as in Figure 2B-2C but with full gene ontology metabolic process category designation or curated category designation. Adjusted p-value $p \leq 0.05$ and fold-change $\geq$ +/-1.5 was considered significant. General categories in legend refer to gene ontology metabolic processes. 'ToxR reg.': ToxR regulon, 'cell comm.': cell communication, 'hyp': hypothetical.

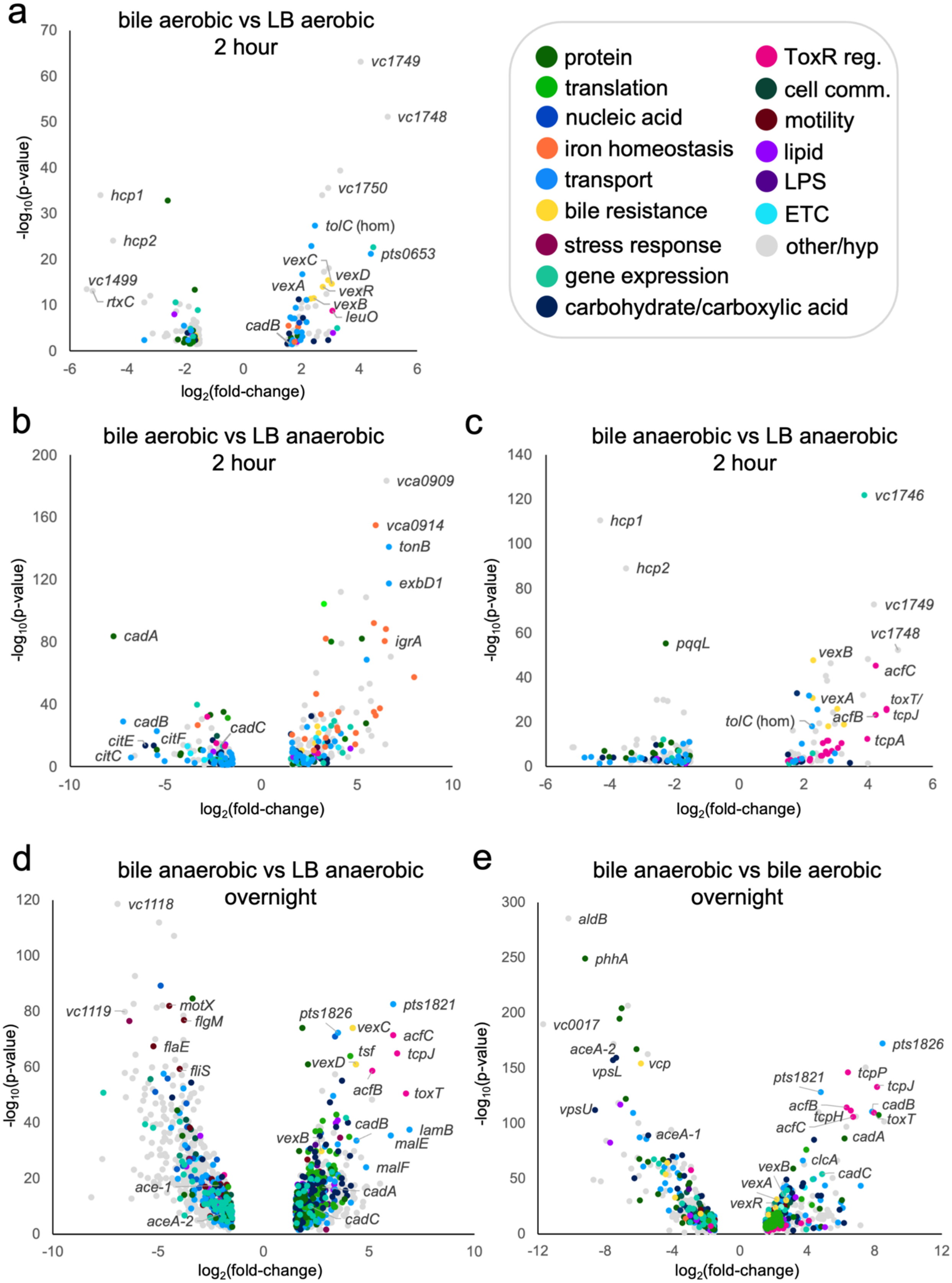
a
bile aerobic vs LB aerobic
2 hour
vc1749
vc1748
vc1750
hcp1
hcp2
tolC (hom)
pts0653
vexC
vexD
vexA
vexR
vexB
leuO
vc1499
rtxC
cadB
-log10(p-value)
log2(fold-change)
protein
translation
nucleic acid
iron homeostasis
transport
bile resistance
stress response
gene expression
carbohydrate/carboxylic acid
ToxR reg.
cell comm.
motility
lipid
LPS
ETC
other/hyp
b
bile aerobic vs LB anaerobic
2 hour
vca0909
vca0914
tonB
exbD1
cadA
igrA
cadB
cadC
citE
citF
citC
c
bile anaerobic vs LB anaerobic
2 hour
vc1746
hcp1
hcp2
vc1749
vc1748
pqqL
vexB
acfC
toxT/
tcpJ
vexA
tolC (hom)
acfB
tcpA
d
bile anaerobic vs LB anaerobic
overnight
vc1118
vc1119
motX
flgM
flaE
fliS
pts1826
pts1821
vexC
acfC
tsf
tcpJ
vexD
toxT
acfB
cadB
lamB
vexB
malE
malF
ace-1
cadA
aceA-2
cadC
e
bile anaerobic vs bile aerobic
overnight
aldB
phhA
vc0017
aceA-2
vcp
vpsL
pts1826
pts1821
tcpP
tcpJ
acfB
cadB
tcpH
toxT
acfC
vpsU
aceA-1
cadA
vexB
clcA
vexA
cadC
vexR

**Supplementary Figure S3**

(**a**) Volcano plot comparing differentially expressed genes in aerobic bile vs. aerobic LB at two hours. (**b**) Volcano plot comparing differentially expressed genes in aerobic bile vs. anaerobic LB at two hours. (**c**) Volcano plot comparing differentially expressed genes in anaerobic bile vs. anaerobic LB at two hours. (**d**) Volcano plot comparing differentially expressed genes in anaerobic bile vs. anaerobic LB overnight cultures. (**e**) Volcano plot comparing differentially expressed genes in anaerobic bile vs. aerobic bile overnight cultures.

For all plots, adjusted p-value $p \leq 0.05$ and fold-change $\geq +/-1.5$ was considered significant. General categories in legend refer to gene ontology metabolic processes. 'ToxR reg.': ToxR regulon, 'cell comm.': cell communication, 'hyp': hypothetical.

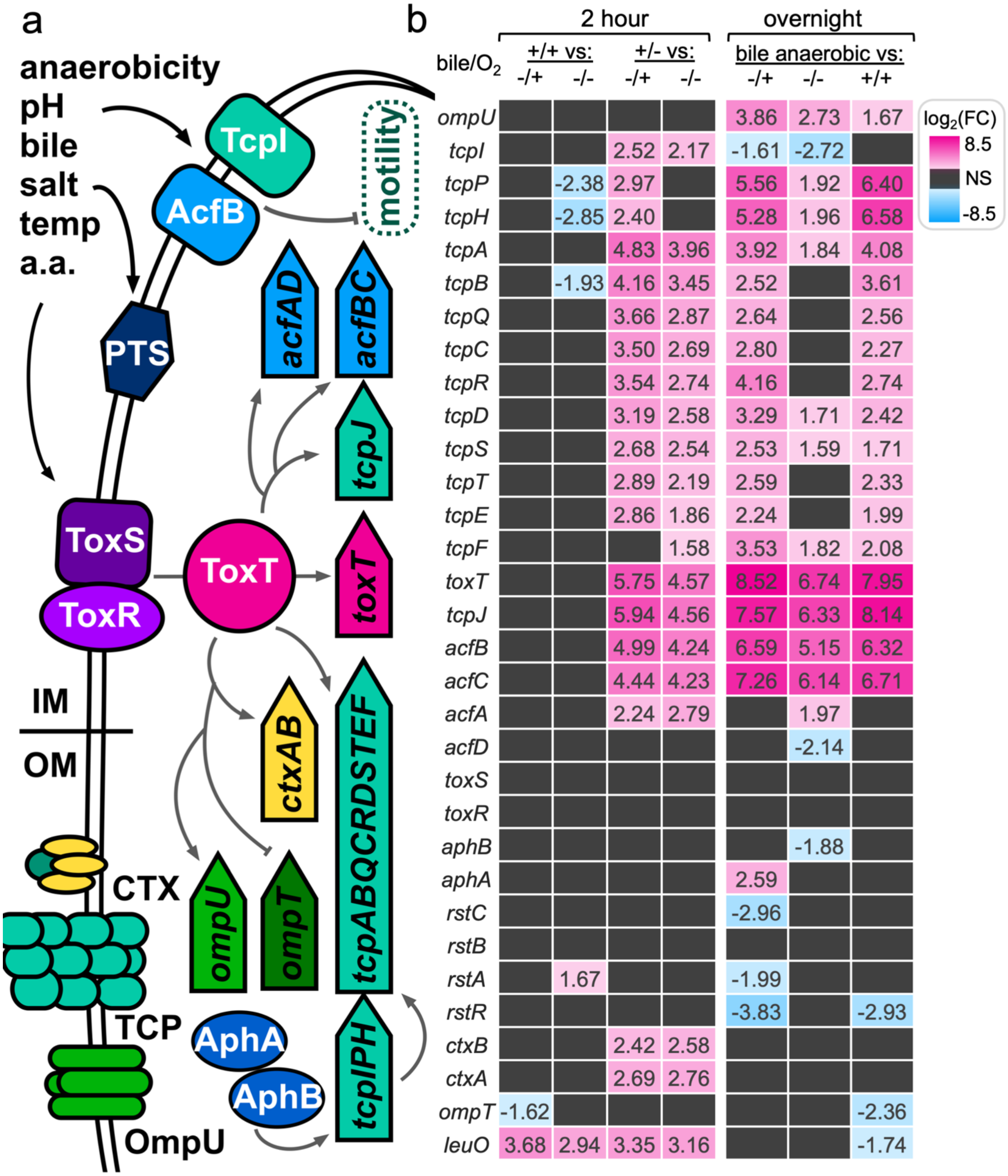


**Supplementary Figure S4**

(**a**) Schematic of select components of the *V. cholerae* toxR virulence regulon. Signals including anaerobicity, pH, bile, osmolarity, temperature ("temp."), and amino acids ("a.a.") are sensed by the ToxRS complex in the inner membrane ("IM"). Stimulation of ToxRS activates expression of *toxT*, which encodes the ToxT transcriptional activator. ToxT activates expression of *ctx* genes encoding the components of cholera toxin and *tcp* genes responsible for building the toxin-

(*Supp. Figure 4 cont.*) coregulated pilus (TCP) at the outer membrane ("OM"), as well as *acf* accessory colonization factor genes involved in a variety of pathogenesis related functions including repression of motility. The ToxR regulon also alters the composition of outer membrane porins by upregulating expression of *ompU* and downregulating *ompT*. Anaerobicity activates AphAB via a thiol-based switch mechanism. AphB is a transcriptional regulator that independently activates *tcpPH* expression. The PTS system senses environmental glucose availability and alters expression of transporters for alternative carbon sources. (refs. 12, 13) (**b**) Heatmap of $log_2$(fold-change) for genes in the ToxR regulon for two-hour and overnight culture comparisons. $log_2$(fold-change) values are listed in individual cells. Legend gives colorimetric approximation of fold-change, NS (black) denotes nonsignificant.

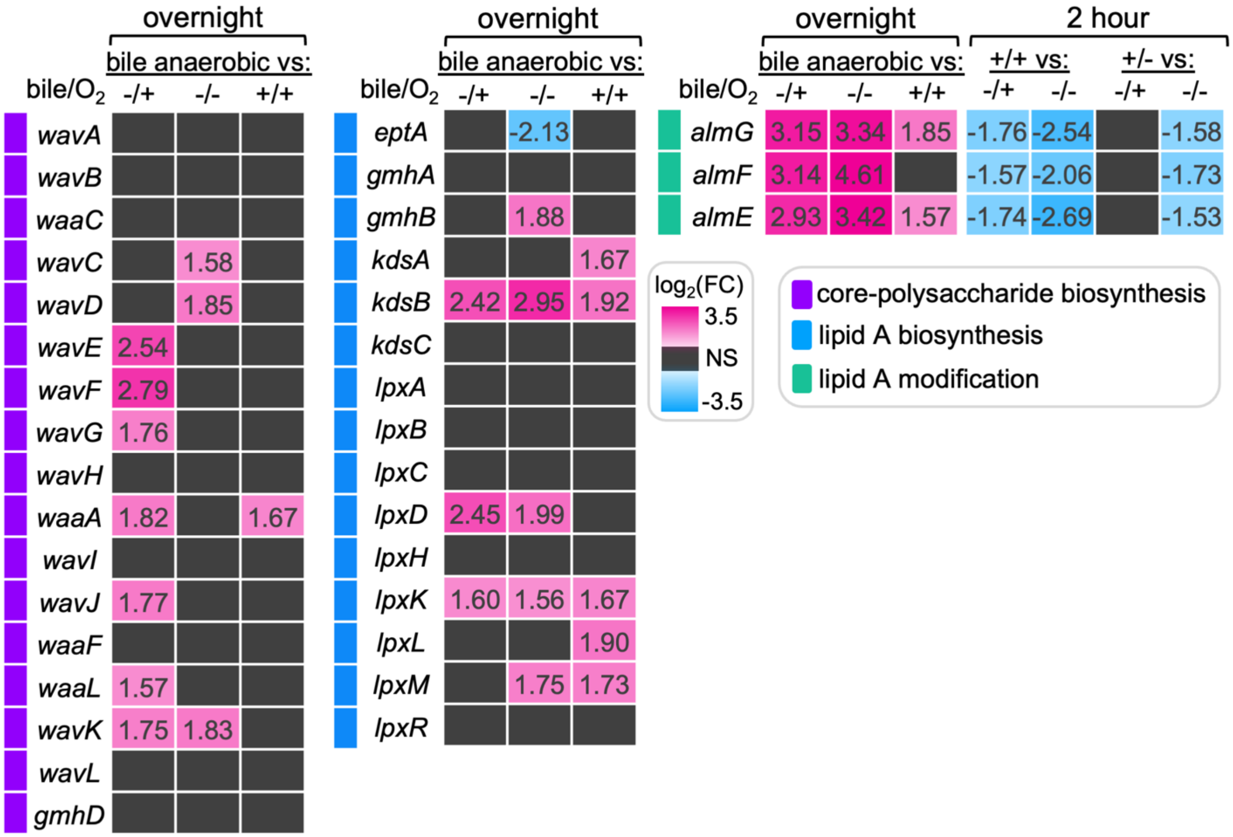


**Supplementary Figure S5**

Heatmap of $log_2$(fold-change) for genes involved in other LPS biosynthetic processes besides O1-antigen biosynthesis (categories on left) for overnight culture comparisons, and additionally at the two-hour timepoint for lipid A modification genes. $log_2$(fold-change) values are listed in individual cells and legend gives colorimetric approximation of fold-change, NS (black) denotes nonsignificant. Second legend indicates gene pathway. Two-hour lipid A/core-polysaccharide biosynthesis gene comparisons and LPS export genes (*lpt* system) were excluded because they were not differentially expressed.

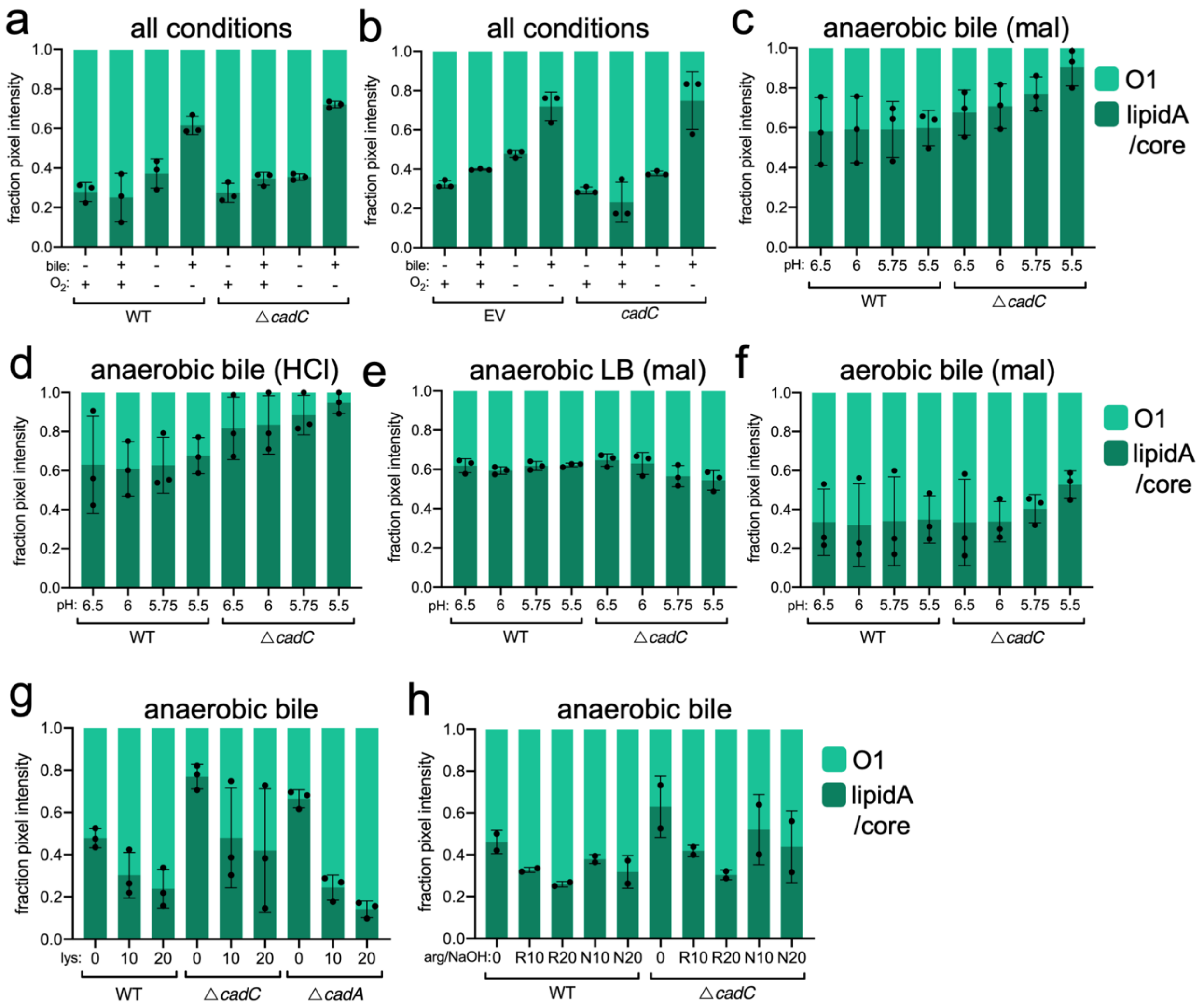


**Supplementary Figure S6**

Pixel intensity quantifications for replicate purified LPS silver stain gels (O1 = light green, lipid A/core = dark green) for the following experiments: (**a**) wild-type (WT) and Δ*cadC V. cholerae* grown overnight in all combinations of culture conditions. (**b**) *V. cholerae* expressing an empty vector (EV) or *cadC* from the *lacZ* locus grown overnight in all combinations of culture conditions. (**c**) wild-type (WT) and Δ*cadC V. cholerae* grown overnight in anaerobic bile culture at low pH (reduced with malic acid). (**d**) wild-type (WT) and Δ*cadC V. cholerae* grown overnight in anaerobic bile culture at low pH (reduced with hydrochloric acid). (**e**) wild-type (WT) and Δ*cadC V. cholerae* grown overnight in anaerobic LB culture at low pH (reduced with malic acid). (**f**) wild-type (WT) and Δ*cadC V. cholerae* grown overnight in aerobic bile culture at low pH (reduced with malic acid). (**g**) wild-type (WT), Δ*cadC*, and Δ*cadA V. cholerae* grown overnight in anaerobic bile culture supplemented with L-lysine (lys, mM). (**h**) wild-type (WT), Δ*cadC*, and Δ*cadA V. cholerae* grown overnight in anaerobic bile culture supplemented with L-arginine (arg/R, mM) or pH adjusted with sodium hydroxide (NaOH/N).

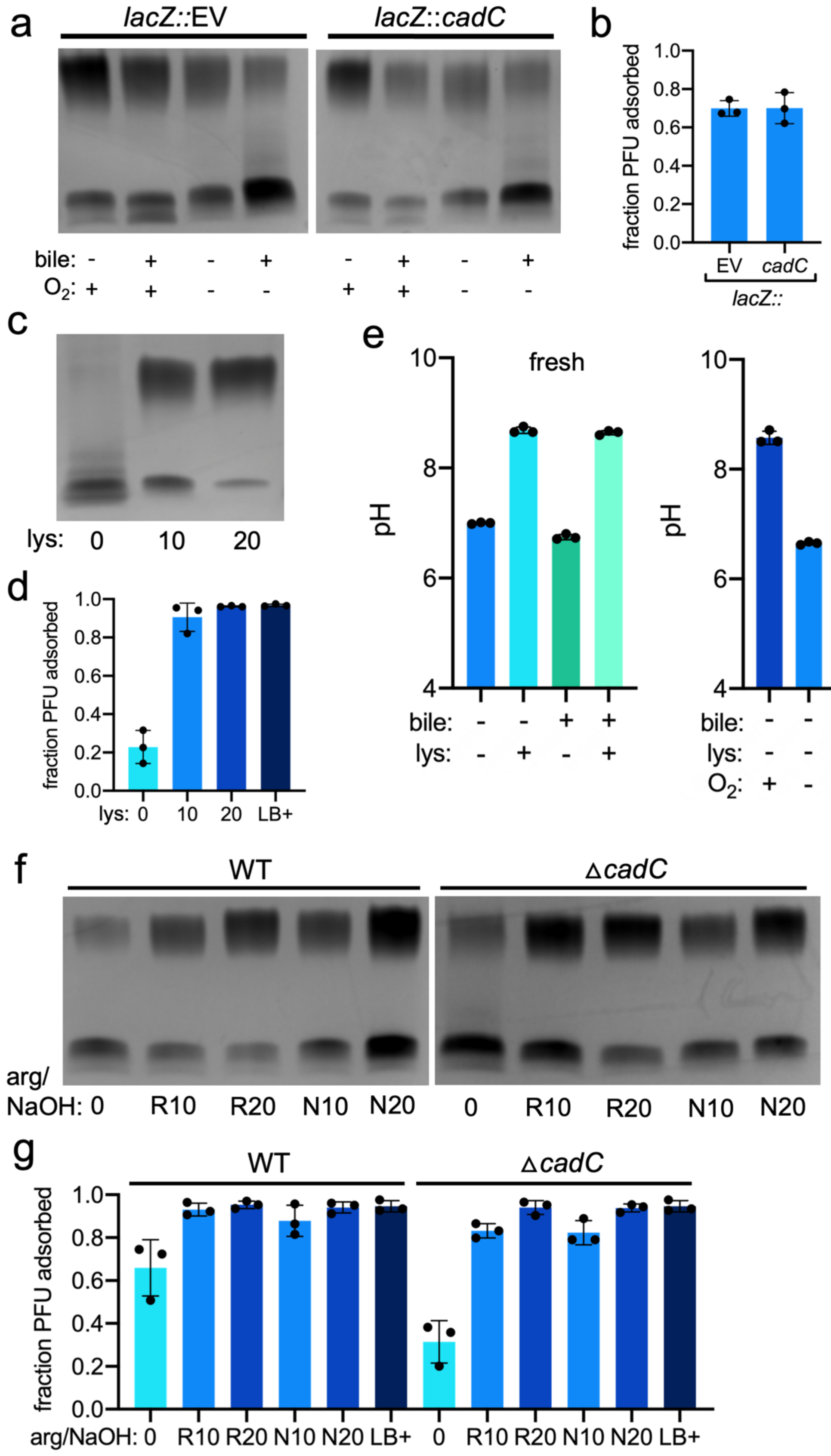

a
lacZ::EV
lacZ::cadC
bile: - + - + - + - +
O2: + + - - + + - -
b
fraction PFU adsorbed
1.0
0.8
0.6
0.4
0.2
0.0
EV
cadC
lacZ::
c
lys: 0 10 20
d
fraction PFU adsorbed
lys: 0 10 20 LB+
e
fresh
spent
pH
10
8
6
4
bile:
lys:
O2:
f
WT
ΔcadC
arg/
NaOH: 0 R10 R20 N10 N20 0 R10 R20 N10 N20
g
WT
ΔcadC
fraction PFU adsorbed
arg/NaOH: 0 R10 R20 N10 N20 LB+ 0 R10 R20 N10 N20 LB+

**Supplementary Figure S7**

(**a**) Purified LPS silver stain of *V. cholerae* grown overnight in all combinations of culture conditions synthetically induced to express *cadC* or an empty vector (EV) in the *lacZ* locus. (**b**) Fraction of ICP1 adsorbed to *V. cholerae* grown overnight in anaerobic bile culture, synthetically induced to express *cadC* or an empty vector (EV) in the *lacZ* locus. (**c**) Purified LPS silver stain of *V. cholerae* Δ*cadA* grown overnight in anaerobic bile culture supplemented with L-lysine (lys, mM). (**d**) Fraction of ICP1 adsorbed to *V. cholerae* Δ*cadA* grown overnight in anaerobic bile culture supplemented with lysine (lys, mM) or aerobic LB (LB+). (**e**) pH measurements of fresh (left) and aerobically/anaerobically spent (right, $O_2$ +/-) LB media with and without 0.5% bile acid supplementation and 20mM L-lysine supplementation. (**f**) Purified LPS silver stain of *V. cholerae* grown overnight in anaerobic bile culture conditions supplemented with L-arginine (arg/R, mM) or pH adjusted with sodium hydroxide to lysine/arginine equivalent (NaOH/N). (**g**) Fraction of ICP1 adsorbed to *V. cholerae* grown overnight in anaerobic bile culture conditions supplemented with L-arginine (arg/R, mM) or pH adjusted with sodium hydroxide to lysine/arginine equivalent (NaOH/N). LB+ denotes aerobic LB control.

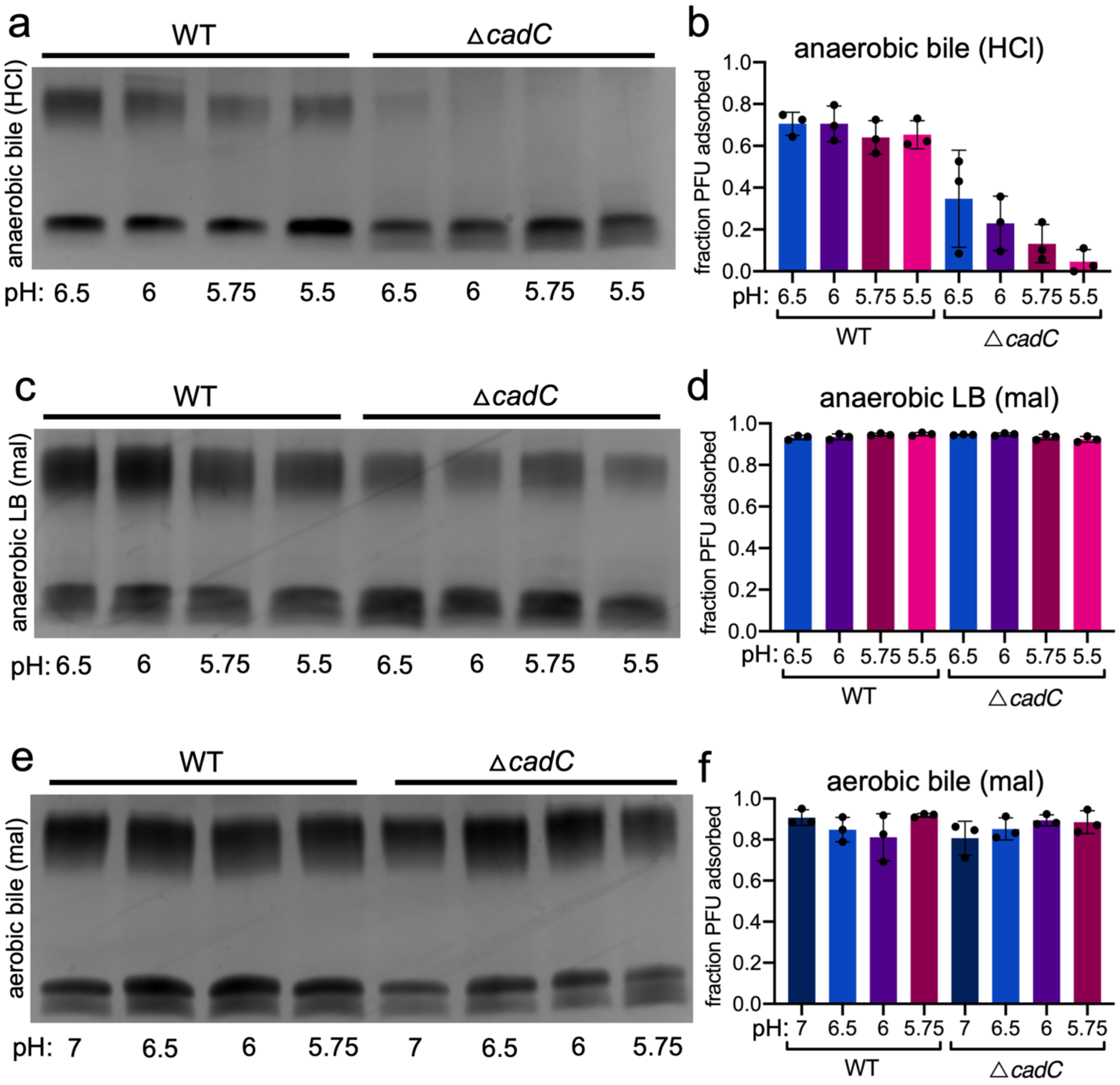


**Supplementary Figure S8**

(**a**) Purified LPS silver stain of *V. cholerae* grown overnight in anaerobic bile culture conditions, pH adjusted with hydrochloric acid. (**b**) Fraction of ICP1 adsorbed to *V. cholerae* grown overnight in anaerobic bile culture conditions, pH reduced with hydrochloric acid (HCl). (**c**) Purified LPS silver stain of *V. cholerae* grown overnight in anaerobic culture conditions, pH reduced with malic acid (mal). (**d**) Fraction of ICP1 adsorbed to *V. cholerae* grown overnight in anaerobic culture conditions, pH reduced with malic acid (mal). (**e**) Purified LPS silver stain of *V. cholerae* grown overnight in aerobic bile culture conditions, pH reduced with malic acid (mal). (**f**) Fraction of ICP1 adsorbed to *V. cholerae* grown overnight in aerobic bile culture conditions, pH reduced with malic acid (mal).

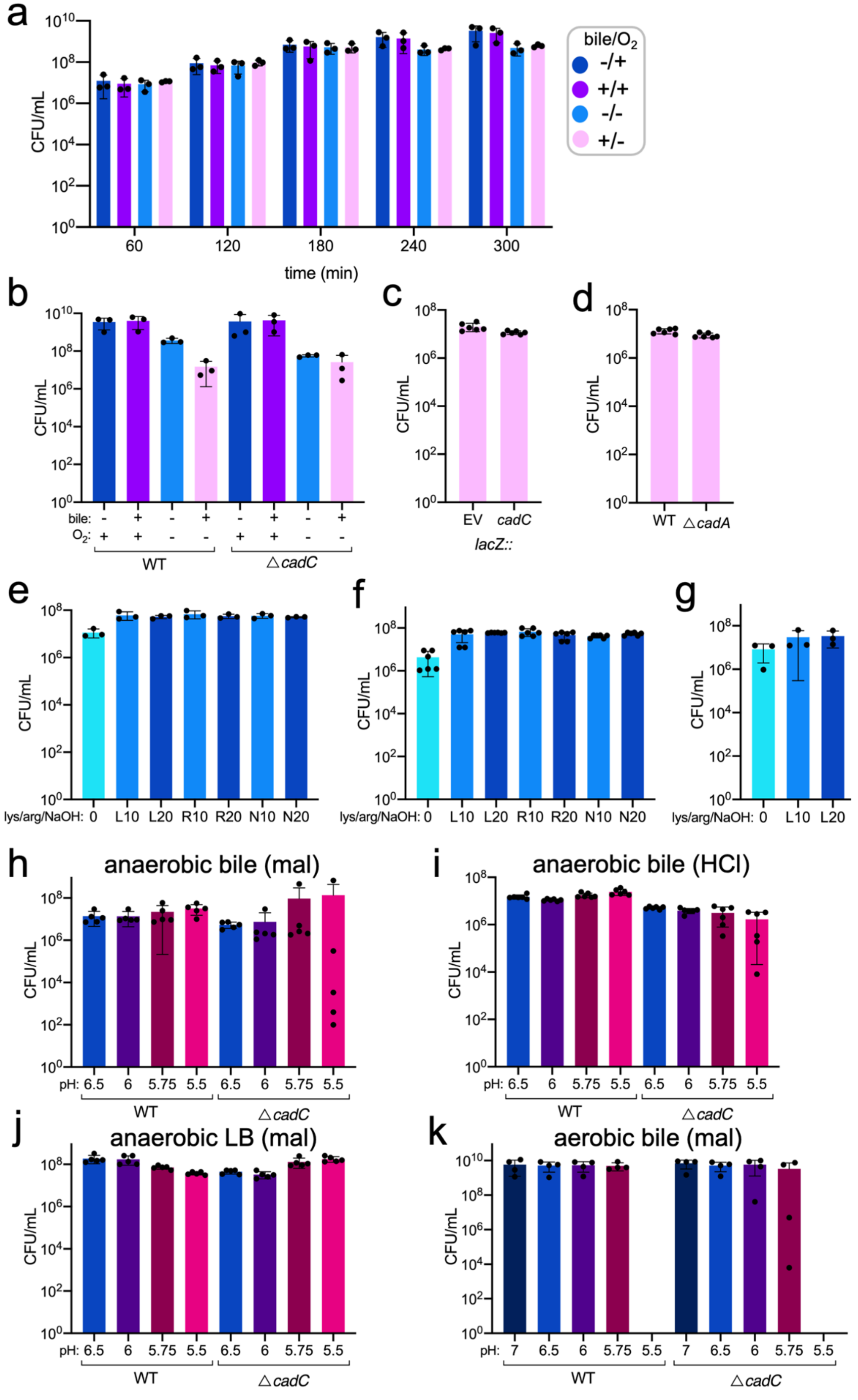

a
CFU/mL
time (min)
60
120
180
240
300
bile/$O_2$
-/+
+/+
-/-
+/-
b
bile:
$O_2$:
WT
△cadC
c
EV
cadC
lacZ::
d
WT
△cadA
e
f
g
lys/arg/NaOH: 0 L10 L20 R10 R20 N10 N20
h
anaerobic bile (mal)
i
anaerobic bile (HCl)
pH: 6.5 6 5.75 5.5 6.5 6 5.75 5.5
j
anaerobic LB (mal)
k
aerobic bile (mal)
pH: 7 6.5 6 5.75 5.5 7 6.5 6 5.75 5.5

**Supplementary Figure S9**

Colony forming units (CFU) quantified for the following experiments:

(a) wild-type *V. cholerae* grown aerobically or anaerobically with or without bile acid supplementation at the indicated time post-inoculation (in minutes). (b) wild-type (WT) and Δ*cadC V. cholerae* grown overnight aerobically and anaerobically ($O_2$+/-) in LB or LB supplemented with 0.5% bile (bile+/-). (c) *V. cholerae* grown overnight in anaerobic bile culture induced to express an empty vector (EV) or *cadC* from the chromosomal *lacZ* locus. (d) wild type (WT) and Δ*cadA V. cholerae* grown overnight in anaerobic bile culture. (e) wild-type *V. cholerae* grown overnight in anaerobic bile culture with lysine (lys/L), arginine (arg/R), or sodium hydroxide (NaOH/N) supplementation. The number next to the letter denotes mM concentration for lysine and arginine, pH matched to mM amino acid concentrations for sodium hydroxide. (f) *V. cholerae* Δ*cadC* grown overnight in anaerobic bile culture with lysine (lys/L), arginine (arg/R), or sodium hydroxide (NaOH/N) supplementation. The number next to the letter denotes mM concentration for lysine and arginine, pH matched to mM amino acid concentrations for sodium hydroxide. (g) *V. cholerae* Δ*cadA* grown overnight in anaerobic bile culture with lysine (lys/L). The number next to the letter denotes mM concentration. (h) Wild-type (WT) and and Δ*cadC V. cholerae* grown overnight in anaerobic bile culture, pH reduced with malic acid. (i) wild-type (WT) and and Δ*cadC V. cholerae* grown overnight in anaerobic bile culture, pH reduced with hydrochloric acid. (j) wild-type (WT) and and Δ*cadC V. cholerae* grown overnight in anaerobic LB culture, pH reduced with malic acid. (k) wild-type (WT) and and Δ*cadC V. cholerae* grown overnight in aerobic bile culture, pH reduced with malic acid.

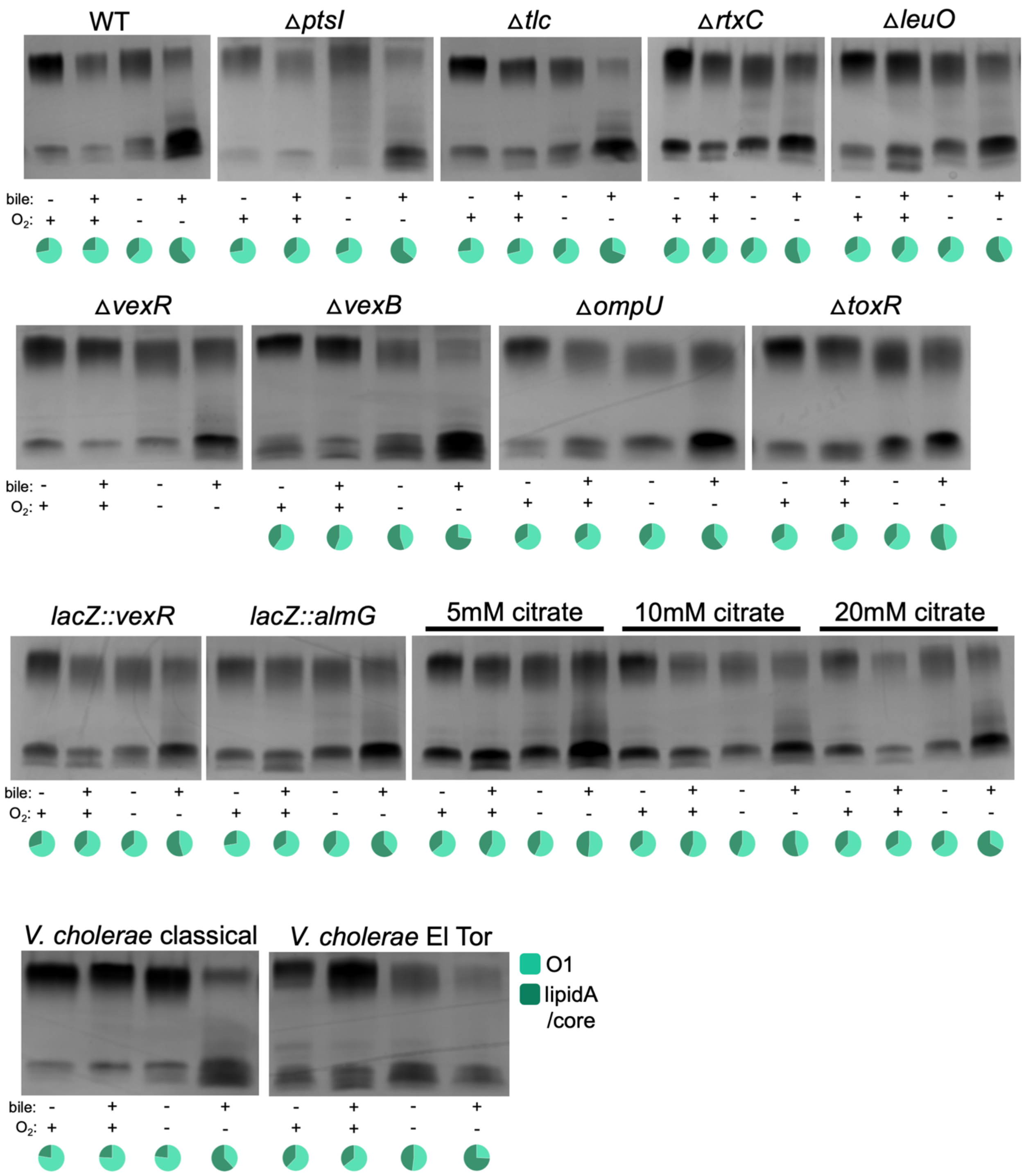


**Supplementary Figure S10**

Representative silver stain gels of LPS purified from *V. cholerae* with indicated genotype (WT: wild type) grown overnight in LB or LB supplemented with 0.5% bile (bile +/-), aerobically or anaerobically ($O_2$ +/-). Gene references are provided in Supplementary Table 2. Gel labeled with citrate represents wild-type *V. cholerae* grown in the indicated concentration of sodium citrate (mM). Pie charts underneath gel images represent pixel intensity quantification (n≥2) of the resulting fraction of intensity from lipid A/core (dark green) and O1-antigen (light green). Quantification was excluded for experiments where only one replicate was conducted (n=1). "*V. cholerae* classical" is a cholera patient isolate from Egypt (1949). "*V. cholerae* clinical" is a cholera patient isolate from Bangladesh (2011). The strain list in Supplementary Table 1 contains reference and source information for all strains.

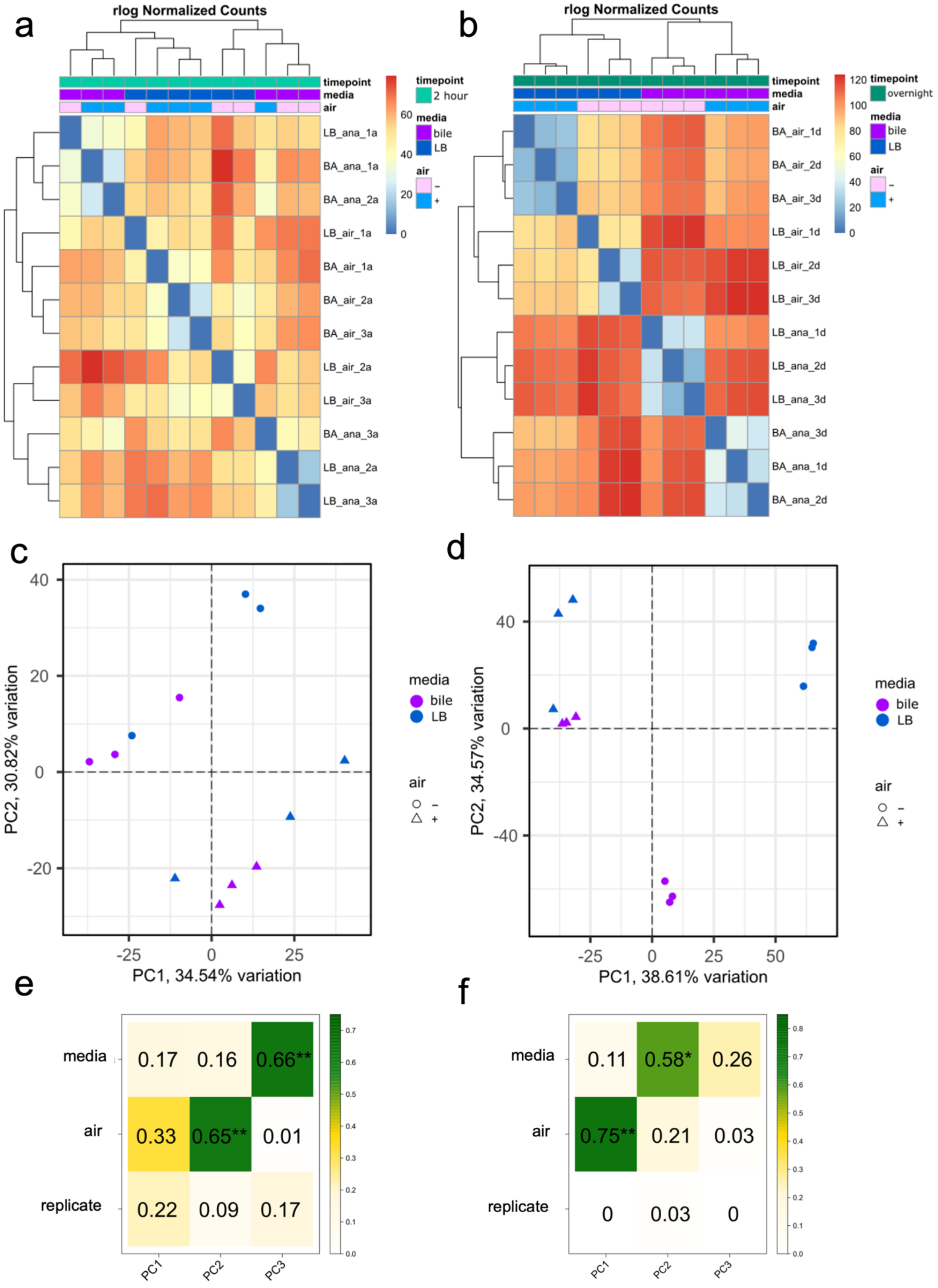

a
rlog Normalized Counts
timepoint
media
air
2 hour
bile
LB
LB_ana_1a
BA_ana_1a
BA_ana_2a
LB_air_1a
BA_air_1a
BA_air_2a
BA_air_3a
LB_air_2a
LB_air_3a
BA_ana_3a
LB_ana_2a
LB_ana_3a
b
rlog Normalized Counts
overnight
BA_air_1d
BA_air_2d
BA_air_3d
LB_air_1d
LB_air_2d
LB_air_3d
LB_ana_1d
LB_ana_2d
LB_ana_3d
BA_ana_3d
BA_ana_1d
BA_ana_2d
c
PC2, 30.82% variation
PC1, 34.54% variation
d
PC2, 34.57% variation
PC1, 38.61% variation
media
bile
LB
air
e
media
air
replicate
0.17 0.16 0.66**
0.33 0.65** 0.01
0.22 0.09 0.17
PC1 PC2 PC3
f
0.11 0.58* 0.26
0.75** 0.21 0.03
0 0.03 0

**Supplementary Figure S11**

Quality assurance analysis of the RNA sequencing individual replicates

(**a**) Two-hour and (**b**) overnight timepoint Euclidian distance matrix, representing the similarity of the individual biological replicates based on the regular logarithmic transformation method. Samples are clustered according to distance and the heatmap scale shows the relative distance between clusters, with lower values representing more similar clusters. (**c**) Two-hour and (**d**) overnight timepoint PCA biplots, showing the comparison of principal components 1 and 2 for each timepoint of samples. Samples are colored according to presence (purple) or absence (blue) of bile in the media conditions and shapes are according to the presence (triangle) or absence (circle) of oxygen during culture. Percent variance described by each principal component is plotted on the x and y axis. (**e**) Two-hour and (**f**) overnight timepoint Eigencor plots, showing the impact of media, aerobicity, and replicates on the variance described by each principal component. The plotted values represent the Pearson $R^2$ coefficient of each treatment across a given principal component. The number of principal components plotted was chosen according to Horn's method in the PCAtools R package.

**Supplementary Table 1.** Strains used in this study
*Reference numbers refer to reference list in text. References with roman numerals in parenthesis are additional references listed below the tables

| Name in text | Strain | Description | Source* |
|---|---|---|---|
| WT | KDS6 | *V. cholerae* E7946. O1, El Tor Ogawa, CTX(+), streptomycin resistant | Lab collection |
| ICP1 | | ICP1_2006_E ΔCRISPR ΔCas2-3 | (I) |
| ICP2 | | | Lab collection |
| Δ*wbeL* | KS201 | *V. cholerae* E7946, *wbeL* open reading frame deleted | 24 |
| *wbeE*-3xFLAG | ZN398 | *V. cholerae* E7946, *vc0244 (wbeE)* tagged with a C-terminal 3x-FLAG tag in the native locus. Also contains a kanamycin resistance cassette in the *lacZ* locus | This study |
| *wbeU*-3xFLAG | ZN403 | *V. cholerae* E7946, *vc0259 (wbeU)* tagged with a C-terminal 3x-FLAG tag in the native locus. Also contains a kanamycin resistance cassette in the *lacZ* locus | This study |
| Δ*cadC* | ZN350 | *V. cholerae* E7946, *cadC* gene replaced with a spectinomycin resistance cassette flanked by FRT sites | This study |
| Δ*cadA* | ZN356 | *V. cholerae* E7946, *cadA* gene replaced with a spectinomycin resistance cassette flanked by FRT sites | This study |
| Δ*ptsI* | ZN329 | *V. cholerae* E7946, *ptsI* gene replaced with a spectinomycin resistance cassette flanked by FRT sites | This study |
| Δ*tlc* | ZN326 | *V. cholerae* E7946, *tlc* genes replaced with a spectinomycin resistance cassette flanked by FRT sites | This study |
| Δ*rtxC* | ZN341 | *V. cholerae* E7946, *rtxC* gene replaced with a spectinomycin resistance cassette flanked by FRT sites | This study |
| Δ*leuO* | ZN344 | *V. cholerae* E7946, *leuO* gene replaced with a spectinomycin resistance cassette flanked by FRT sites | This study |
| Δ*vexR* | ZN347 | *V. cholerae* E7946, *vexR* gene replaced with a spectinomycin resistance cassette flanked by FRT sites | This study |
| Δ*vexB* | ZN57 | *V. cholerae* E7946, *vexB* gene replaced with a spectinomycin resistance cassette flanked by FRT sites | This study |
| Δ*ompU* | KS799 | *V. cholerae* E7946, *ompU* open reading frame deleted | (II) |

| | | | |
|---|---|---|---|
| Δ*toxR* | KS801 | *V. cholerae* E7946, *toxR* open reading frame deleted | (III) |
| *lacZ::*EV | KS1500 | *V. cholerae* E7946, *lacZ* gene replaced with an empty cassette under IPTG-inducible ($P_{tac}$) and theophylline-inducible (RiboE) promoters and a kanamycin resistance marker | (I) |
| *lacZ::vexR* | ZN332 | *V. cholerae* E7946, *lacZ* gene replaced with *vexR* under IPTG-inducible ($P_{tac}$) and theophylline-inducible (RiboE) promoters and a kanamycin resistance marker | This study |
| *lacZ::almG* | ZN335 | *V. cholerae* E7946, *lacZ* gene replaced with *almG* under IPTG-inducible ($P_{tac}$) and theophylline-inducible (RiboE) promoters and a kanamycin resistance marker | This study |
| *lacZ::cadC* | ZN338 | *V. cholerae* E7946, *lacZ* gene replaced with *cadC* under IPTG-inducible ($P_{tac}$) and theophylline-inducible (RiboE) promoters and a kanamycin resistance marker | This study |
| *V. cholerae* classical | KS808 | *V. cholerae* clinical patient isolate from Egypt, 1949 | (IV) |
| *V. cholerae* El Tor | KDS1 | *V. cholerae* clinical patient isolate from Bangladesh, 2011 | (V) |

**Supplementary Table 2.** Gene names references in text
*Reference numbers refer to reference list in text. References with roman numerals in parenthesis are additional references listed below the tables

| name (text) | VCID | locus tag | annotation | description | ref* |
|---|---|---|---|---|---|
| *vexA* | VC0165 | CSW01_ RS00860 | hypothetical protein | RND bile efflux component | 33 |
| *vexB* | VC0164 | CSW01_ RS00855 | multidrug resistance protein | RND bile efflux component | 33 |
| *vexC* | VC1756 | CSW01_ RS08835 | periplasmic linker protein | RND bile efflux component | 33 |
| *vexD* | VC1757 | CSW01_ RS08840 | AcrB/AcrD/AcrF family transporter | RND bile efflux component | 33 |
| *ompU* | VC0633 | CSW01_ RS03335 | porin | outer membrane porin involved in bile resistance | 30 |
| *tcpI* | VC0825 | CSW01_ RS04260 | toxin co-regulated pilus biosynthesis protein I | TCP methyl-accepting chemoreceptor | 36 |

| | | | | | |
|---|---|---|---|---|---|
| *tcpP* | VC0826 | CSW01_ RS04265 | toxin co-regulated pilus biosynthesis protein P | TCP putative regulatory protein | 36 |
| *tcpH* | VC0827 | CSW01_ RS04270 | toxin co-regulated pilus biosynthesis protein H | | 36 |
| *tcpA* | VC0828 | CSW01_ RS04275 | toxin co-regulated pilin | TCP major subunit | 36 |
| *tcpB* | VC0829 | CSW01_ RS04280 | toxin co-regulated pilus biosynthesis protein B | TCP minor subunit | 36 |
| *tcpQ* | VC0830 | CSW01_ RS04285 | toxin co-regulated pilus biosynthesis protein Q | | 36 |
| *tcpC* | VC0831 | CSW01_ RS04290 | toxin co-regulated pilus biosynthesis outer membrane protein C | TCP outer membrane lipoprotein | 36 |
| *tcpR* | VC0832 | CSW01_ RS04295 | toxin co-regulated pilus biosynthesis protein R | | 36 |
| *tcpD* | VC0833 | CSW01_ RS04300 | toxin co-regulated pilus biosynthesis protein D | | 36 |
| *tcpS* | VC0834 | CSW01_ RS04305 | toxin co-regulated pilus biosynthesis protein S | | 36 |
| *tcpT* | VC0835 | CSW01_ RS04310 | toxin co-regulated pilus biosynthesis protein T | TCP membrane-associated ATPase | 36 |
| *tcpE* | VC0836 | CSW01_ RS04315 | toxin co-regulated pilus biosynthesis protein E | | 36 |
| *tcpF* | VC0837 | CSW01_ RS04320 | toxin co-regulated pilus biosynthesis protein F | TCP putative outer membrane channel | 36 |
| *toxT* | VC0838 | CSW01_ RS04325 | TCP pilus virulence regulatory protein | regulatory protein | 35 |
| *tcpJ* | VC0839 | CSW01_ RS04330 | leader peptidase TcpJ | signal peptidase | 36 |
| *acfB* | VC0840 | CSW01_ RS04335 | accessory colonization factor AcfB | accessory colonization factor | 38 |
| *acfC* | VC0841 | CSW01_ RS04340 | accessory colonization factor AcfC | accessory colonization factor | 39 |
| *acfA* | VC0844 | CSW01_ RS04355 | accessory colonization factor AcfA | accessory colonization factor | 37 |
| *acfD* | VC0845 | CSW01_ RS04360 | accessory colonization factor AcfD | accessory colonization factor | (VI) |
| *toxS* | VC0983 | CSW01_ RS05035 | regulatory protein ToxS | | 35 |
| *toxR* | VC0984 | CSW01_ RS05040 | cholera toxin transcriptional activator | | 35 |
| *aphB* | VC1049 | CSW01_ RS05355 | LysR family transcriptional regulator | accessory virulence factor | 84 |
| *aphA* | VC1050 | CSW01_ RS05360 | response regulator | accessory virulence factor | 85 |

| | | | | | |
|---|---|---|---|---|---|
| *rstC* | VC1452 | CSW01_ RS07300 | RstC protein | | 41 |
| *rstB* | VC1453 | CSW01_ RS07305 | RstB1 protein | | 41 |
| *rstA* | VC1463 | CSW01_ RS07310 | RstA2 protein | | 41 |
| *rstR* | VC1464 | CSW01_ RS07315 | transcriptional repressor RstR | | 41 |
| *ctxB* | VC1456 | CSW01_ RS07325 | cholera enterotoxin subunit B | | 40 |
| *ctxA* | VC1457 | CSW01_ RS07330 | cholera enterotoxin subunit A | | 40 |
| *ompT* | VC1854 | CSW01_ RS09310 | porin | outer membrane porin | 30 |
| *leuO* | VC2485 | CSW01_ RS12615 | leucine transcriptional activator | | (VII) |
| *almE* | VC1577 | CSW01_ RS07935 | enterobactin synthetase subunit F | lipid A modification amino acid ligase | 45 |
| *almF* | VC1578 | CSW01_ RS07940 | hypothetical protein | lipid A modification glycine carrier protein | 45 |
| *almG* | VC1579 | CSW01_ RS07945 | hypothetical protein | lipid A modification glycine transferase | 45 |
| *vexR* | VC0166 | CSW01_ RS00865 | TetR family transcriptional regulator | bile RND efflux regulator | 33 |
| *ptsI* | VC0965 | CSW01_ RS04940 | phosphoenolpyruvate-protein phosphotransferase EI | | 51 |
| *tlc* | VC1466-70, VC1472-6 | CSW01_ RS07380 -405, CSW01_ RS07415 -45 | | Satellite phage of CTX (2 copies) | (VIII) |
| *cadC* | VC0278 | CSW01_ RS01450 | DNA-binding transcriptional activator CadC | weak acid tolerance system | 54 |
| *cadB* | VC0280 | CSW01_ RS01455 | lysine/cadaverine antiporter | weak acid tolerance system | 54 |
| *cadA* | VC0281 | CSW01_ RS01460 | lysine decarboxylase, inducible | weak acid tolerance system | 54 |
| *rtxA* | VC1451 | CSW01_ RS07295 | RTX toxin RtxA | | 52 |
| *rtxC* | VC1450 | CSW01_ RS07290 | RTX toxin activating protein | | 52 |
| *clcA* | VCA0526 | CSW01_ RS16970 | chloride channel protein | | 64 |

| | | | | | |
|---|---|---|---|---|---|
| *makD* | VC0880 | CSW01_ RS04530 | hypothetical protein | motility associated killing factor D | 76 |
| *makC* | VC0881 | CSW01_ RS04535 | hypothetical protein | motility associated killing factor C | 76 |
| *makB* | VC0882 | CSW01_ RS04540 | hypothetical protein | motility associated killing factor B | 76 |
| *makA* | VC0884 | CSW01_ RS04545 | acetyltransferase-like protein | motility associated killing factor A | 76 |
| *aceA-1* | VC0734 | CSW01_ RS03835 | malate synthase | | 79 |
| *aceA-2* | VCA0957 | CSW01_ RS19015 | malate synthase | | 79 |
| *glmS* | VC0487 | CSW01_ RS02600 | glucosamine--fructose-6-phosphate aminotransferase | | (IX) |
| *vprA* | VC1320 | CSW01_ 06665 | DNA-binding response regulator aka carR | | 46 |
| *vprB* | VC1319 | CSW01_ RS06680 | sensor histidine kinase aka carS | | 46 |
| *manC* | VC0241 | CSW01_ RS01280 | mannose-1-phosphate guanylyltransferase | O-biosynthetic protein-perosamine | 47 |
| *manB* | VC0242 | CSW01_ RS01285 | phosphomannomutase | O-biosynthetic protein-perosamine | 47 |
| *gmd* | VC0243 | CSW01_ RS01290 | GDP-mannose 4,6-dehydratase | O-biosynthetic protein-perosamine | 47 |
| *wbeE* | VC0244 | CSW01_ RS01295 | perosamine synthase | O-biosynthetic protein-perosamine | 47 |
| *wbeG* | VC0245 | CSW01_ RS01300 | RfbG protein | O-biosynthetic protein-other | 47 |
| *wzm* | VC0246 | CSW01_ RS01305 | lipopolysaccharide/O-antigen transport protein | O-biosynthetic protein-transport | 47 |
| *wzt* | VC0247 | CSW01_ RS01310 | lipopolysaccharide/O-antigen transport protein | O-biosynthetic protein-transport | 47 |
| *wbeK* | VC0248 | CSW01_ RS01315 | acyl carrier protein | O-biosynthetic protein-tetronate | 47 |
| *wbeL* | VC0249 | CSW01_ RS01320 | RfbL protein | O-biosynthetic protein-tetronate | 47 |
| *wbeM* | VC0250 | CSW01_ RS01325 | iron-containing alcohol dehydrogenase | O-biosynthetic protein-tetronate | 47 |
| *wbeN* | VC0251 | CSW01_ RS01330 | acyl protein synthase/acyl-CoA reductase RfbN | O-biosynthetic protein-tetronate | 47 |
| *wbeO* | VC0252 | CSW01_ RS01335 | acetyltransferase | O-biosynthetic protein-tetronate | 47 |

| | | | | | |
|---|---|---|---|---|---|
| *wbeP* | VC0253 | CSW01_ RS01340 | unknown | O-biosynthetic protein-tetronate | 47 |
| IS1358d1 | VCA0493 | CSW01_ RS01345 | IS1004 transposase | | 47 |
| *wbeT* | VC0258 | CSW01_ RS01355 | RfbT-like protein | O-biosynthetic protein-tetronate | 47 |
| *wbeV* | VC0259 | CSW01_ RS01360 | lipopolysaccharide biosynthesis protein RfbV | O-biosynthetic protein-other | 47 |
| *wbeU* | VC0260 | CSW01_ RS01365 | mannosyltransferase | O-biosynthetic protein-other | 47 |
| *galE* | VC0262 | CSW01_ RS01370 | UDP-glucose 4-epimerase | O-biosynthetic protein-other | 47 |
| *wbeW* | VC0263 | CSW01_ RS01375 | galactosyl-transferase | O-biosynthetic protein-other | 47 |
| *wavA* | VC0223 | CSW01_ RS01195 | ADP-heptose--LPS heptosyltransferase II | core-polysaccharide biosynthesis | 47 |
| *wavB* | VC0224 | CSW01_ RS01200 | lipopolysaccharide biosynthesis glycosyltransferase | core-polysaccharide biosynthesis | 47 |
| *waaC* | VC0225 | CSW01_ RS01205 | lipopolysaccharide biosynthesis protein | core-polysaccharide biosynthesis | 47 |
| *wavC* | VC0227 | CSW01_ RS01210 | 3-deoxy-D-manno-octulosonic acid kinase | core-polysaccharide biosynthesis | 47 |
| *wavD* | VC0228 | CSW01_ RS01215 | hypothetical protein | core-polysaccharide biosynthesis | 47 |
| *wavE* | VC0229 | CSW01_ RS01220 | hypothetical protein | core-polysaccharide biosynthesis | 47 |
| *wavF* | VC0230 | CSW01_ RS01225 | hypothetical protein | core-polysaccharide biosynthesis | 47 |
| *wavG* | VC0231 | CSW01_ RS01230 | hypothetical protein | core-polysaccharide biosynthesis | 47 |
| *wavH* | VC0232 | CSW01_ RS01235 | hypothetical protein | core-polysaccharide biosynthesis | 47 |
| *waaA* | VC0233 | CSW01_ RS01240 | 3-deoxy-D-manno-octulosonic acid transferase | core-polysaccharide biosynthesis | 47 |
| *wavI* | VC0234 | CSW01_ RS01245 | hypothetical protein | core-polysaccharide biosynthesis | 47 |
| *wavJ* | VC0235 | CSW01_ RS01250 | lipopolysaccharide biosynthesis protein | core-polysaccharide biosynthesis | 47 |
| *waaF* | VC0236 | CSW01_ RS01255 | ADP-heptose--LPS heptosyltransferase II | core-polysaccharide biosynthesis | 47 |
| *waaL* | VC0237 | CSW01_ RS01260 | hypothetical protein | core-polysaccharide biosynthesis | 47 |

| | | | | | |
|---|---|---|---|---|---|
| *wavK* | VC0238 | CSW01_ RS01265 | hexapaptide repeat-containing transferase | core-polysaccharide biosynthesis | 47 |
| *wavL* | VC0239 | CSW01_ RS01270 | hypothetical protein | core-polysaccharide biosynthesis | 47 |
| *gmhD* | VC0240 | CSW01_ RS01275 | ADP-L-glycero-D-mannoheptose-6-epimerase | core-polysaccharide biosynthesis | 47 |
| *eptA* | VCA1102 | CSW01_ RS19695 | hypothetical protein | lipid A biosynthesis | (X) |
| *gmhA* | VC2230 | CSW01_ RS11395 | phosphoheptose isomerase | lipid A biosynthesis | (X) |
| *gmhB* | VC0908 | CSW01_ RS04660 | D,D-heptose 1,7-bisphosphate phosphatase | lipid A biosynthesis | (X) |
| *kdsA* | VC2175 | CSW01_ RS10855 | 2-dehydro-3-deoxyphosphooctonate aldolase | lipid A biosynthesis | (X) |
| *kdsB* | VC1875 | CSW01_ RS09420 | 3-deoxy-manno-octulosonate cytidylyltransferase | lipid A biosynthesis | (X) |
| *lpxA* | VC2248 | CSW01_ RS11480 | acyl-(acyl-carrier-protein)--UDP-N-acetylglucosamine O-acyltransferase | lipid A biosynthesis | (X) |
| *lpxB* | VC2247 | CSW01_ RS11475 | lipid-A-disaccharide synthase | lipid A biosynthesis | (X) |
| *lpxC* | VC2396 | CSW01_ RS12170 | UDP-3-O-[3-hydroxymyristoyl] N-acetylglucosamine deacetylase | lipid A biosynthesis | (X) |
| *lpxD* | VC2250 | CSW01_ RS11490 | UDP-3-O-[3-hydroxymyristoyl] glucosamine N-acyltransferase | lipid A biosynthesis | (X) |
| *lpxH* | VC1850 | CSW01_ RS09290 | UDP-2,3-diacylglucosamine hydrolase | lipid A biosynthesis | (X) |
| *lpxK* | VC1877 | CSW01_ RS09430 | tetraacyldisaccharide 4'-kinase | lipid A biosynthesis | (X) |
| *lpxL* | VC0213 | CSW01_ RS01140 | lipid A biosynthesis lauroyl acyltransferase | lipid A biosynthesis | (X) |
| *lpxM* | VC0212 | CSW01_ RS01135 | lipid A biosynthesis (KDO)2-(lauroyl)-lipid IVA acyltransferase | lipid A biosynthesis | (X) |
| *lpxR* | VC1867 | CSW01_ RS09380 | hypothetical protein | lipid A biosynthesis | (X) |
