## Appendix for "Adaptation to bile and anaerobicity limits *Vibrio cholerae* phage adsorption"

### Appendix: Extended RNA-sequencing data analysis

#### RNA-sequencing quality assessment by principal component analysis

We performed multiple parallel analyses to confirm that the RNA sequencing samples clustered as expected between samples. First, we transformed the transcript count matrix using the regularized logarithm (rlog) transformation. Using this transformed dataset, we generated a Euclidian distance matrix of the transcript counts at both the two-hour (Supplementary Figure S11A) timepoint and the overnight timepoint (Supplementary Figure S11B). Replicates from the overnight analysis clustered more tightly with each other than those from the two-hour analysis, consistent with these earlier samples still undergoing adaptation to their respective culture conditions. To further assess the quality of our data, we used principal component analysis (PCA) to assess the relatedness of the same transformed count data. PCA of the full dataset revealed clustering of replicates between timepoints but not treatment conditions, informing our further differential expression analysis to be within timepoints. Plotting the replicates on the PC1 and PC2 axes revealed loose clustering of replicates at the two-hour timepoint (Supplementary Figure S11C) and strict clustering of samples at the overnight timepoint (Supplementary Figure S11D). To reveal the contribution of each individual treatment condition to each principal component, we generated Eigencor plots showing the Pearson pairwise $r^2$ correlates between samples at the two-hour (Supplementary Figure S11E) and overnight (Supplementary Figure S11F) timepoints. This analysis further confirmed the robust differences in transcriptional profile captured by PC1 and PC2 at the overnight timepoint to be due to impacts of aerobicity and media conditions, respectively. At the two-hour timepoint, PC2 and PC3 were correlated with aerobicity and media, showing the same trends as the overnight timepoint, despite the decrease in differences between sample treatments, which we attribute to active transcriptional reprogramming to adapt to the culture conditions.

#### RNA-sequencing extended results discussion

RNA sequencing analysis revealed an intersection across all compared conditions of less than 10% of differentially regulated genes at 2 hours (23/411 downregulated and 28/320 upregulated genes, Supplementary Figure S2A) compared to 13% of differentially expressed genes at the overnight timepoint (117/1077 upregulated and 123/1324 downregulated, Supplementary Figure S2B, Supplementary Data Sheet 2). We had hoped to identify a differentially expressed transcriptional regulator, protease, or gene involved with O1-antigen biosynthesis shared across all comparisons in our RNA-seq analysis where we observed reduced O1-antigen decoration. Instead, the pool of overlapping genes did not provide any immediately obvious connections to O1-antigen production. However, the shared differentially expressed genes may contribute to contextualizing the O1-antigen reduction phenotype in *V. cholerae* pathogenesis. Some shared upregulated genes met our expectations: *vexB* bile efflux upregulation in bile correlates with previous descriptions of the system[34], and upregulation of elongation factor G may be indicative of a cellular attempt to achieve homeostasis via global upregulation of protein synthesis.

Across all comparisons, we observed a reduction in the expression of genes involved in two different toxin systems: the multifunctional auto-processing RTX toxin (RTX toxin activator *rtxC* and the hypothetical gene immediately upstream, *vc1499*)[52] and the Mak motility-associated-killing cytotoxin complex (*makDC*, *vca0880* and *vca0881*[76,77]). Since it is expected that these toxins play a role in virulence, it seemed odd that they would be consistently downregulated in anaerobic bile conditions that appear to mimic the intestinal environment. However, RTX toxin has been shown to be important not for the establishment of colonization but instead for the maintenance of long-term colonization, where *V. cholerae* appears to rely on RTX and other accessory virulence factors instead of TCP and CTX[78]. If the overnight anaerobic bile and two-hour bile conditions more closely resemble signals from the intestinal environment prior to *V. cholerae* colonization, then it would make sense that factors required later may be downregulated by these signals and instead respond to signals of an intestine that has already been colonized (e.g. bile replaced with watery diarrhea induced by CTX expression). The case could be similar for the Mak toxin system, especially because it relies on a flagellar secretion system that is transcriptionally downregulated by accessory colonization factors controlled by the ToxR regulon[76,77]. However, the Mak system is conserved among a variety of *Vibrio* species including pathogens of fish and has not yet been explored in a mammalian animal model, so it may also be an important system for survival in the aquatic environment. Taken together, these results support the utilization of isolated environmental signals to more deeply understand the complex regulatory pathways in organisms like *V. cholerae* that oscillate between dramatically different environments.

A few other notable trends in the RNA-seq data further supported our conclusions about limited availability of central carbon metabolism intermediates in cells producing less O1-decorated LPS. For all comparisons at the overnight timepoint, both copies of the *V. cholerae* malate synthase gene (*aceA-1, aceA-2*[79]) were significantly downregulated (Supplementary Figure S2D, S3E). Malate is the precursor for tetronate biosynthesis, a required step in O1-antigen biosynthesis[31]. Transcriptional downregulation of malate synthase would likely negatively impact availability of malate for O1-antigen biosynthesis. *glmS* (an enzyme that catalyzes the first step in the hexosamine biosynthetic pathway[80]) was similarly consistently downregulated in all overnight comparisons (Supplementary Table). The hexosamine pathway redirects fructose 6-phosphate from glycolysis to synthesize amino acids, amino sugars, and nucleic acid sugars necessary for both LPS and peptidoglycan synthesis, meaning downregulation of this pathway would negatively impact the availability of a wide variety of cellular macromolecules, including necessary LPS precursors. We infer that transcriptional downregulation of these biosynthesis enzymes leads to a reduction in the availability of their products, but this remains to be experimentally determined.

We also observed perturbations in expression of the *almEFG* lipid A remodeling system: the system was upregulated in every comparison at two hours and downregulated in every overnight comparison. AlmEFG modify *V. cholerae* lipid A with glycine to confer increased resistance to antimicrobial peptides found in the gut[45]. This system is linked to the ToxR regulon via LeuO-mediated activation of *vprAB*, which activate *almEFG*[46,81]. We were curious if O1-antigen substitution was controlled by a similar *toxR*-dependent process, however when we grew a Δ*toxR* strain in anaerobic bile conditions we observed a reduction in O1-antigen substitution similar to wild-type, indicating that the O1-antigen reduction occurs independent of ToxR regulon activation (Supplementary Figure S10). We also tested mutants in genes immediately downstream of ToxR/ToxT-mediated activation (*leuO* and *ompU*) and observed LPS production similar to wild-type (Supplementary Figure S10), confirming that unlike lipid A remodeling, O1-antigen reduction occurs independent of the ToxR regulon.

The results of our RNA sequencing analysis confirm the expectations established by previous analyses via microarray and 2D gel proteomics examining the effects of bile acids and anaerobicity in *V. cholerae*. Consistent with this work, previous microarray-based characterization of the *V. cholerae* transcriptional response to bile identified transcriptional upregulation of *vexAB* and *vexCD* RND bile efflux systems, as well as global upregulation of transport-associated genes and downregulation of motility genes[34]. The microarray analysis also detected transcriptional downregulation of a subset of virulence factors including *ctxAB* and some TCP components in the presence of bile. This result correlates with other work demonstrating that while ToxR activity is stimulated by bile, the transcriptional activation activity of ToxT may be negatively regulated by bile[82,83] allowing for fine-tuning of virulence transcriptional profiles after ToxR activation. While we observed differential upregulation of the complete suite of virulence genes when examining the anaerobic bile condition, bile alone did not appear to stimulate transcriptional changes in virulence genes (Supplementary Figure S4), consistent with the necessity of both bile and anaerobicity to activate the complete virulence cascade. The isolated effect of anaerobicity has been previously examined via 2D gel proteomics[60] and identified several trends also represented in the RNA-seq analysis in this work. Iron homeostasis and vibriobactin biosynthetic enzymes were observed to be less abundant in anaerobic conditions, consistent with our observations of decreased transcriptional activity of the cognate genes. Proteomics also identified anaerobicity-driven increases in abundance of accessory colonization factors and TCP components but not CTX, supporting the hypothesis that the ToxR regulon is not an all-on or all-off switch but instead interprets and responds to different environmental signals by altering expression of different components downstream of ToxR. This also contextualizes our observations of differential expression of these virulence factors in the overnight condition, where we detect weaker differential expression of TCP and accessory colonization factors comparing the anaerobic bile condition to anaerobic LB, while the comparisons to aerobic conditions demonstrated more robust transcriptional upregulation of these virulence components (Supplementary Figure S3). While we cannot draw direct comparisons between transcriptomic and proteomic/microarray data sets, a general understanding of the global effects of anaerobicity and bile individually aided in contextualizing the variable differential expression between condition comparisons in our transcriptomic data set and supports the existing understanding of the complex nature of *V. cholerae* virulence gene regulation.

One well-characterized component of the ToxR regulon that is post-transcriptionally regulated by anaerobicity is transcription factor AphB[84], which (together with AphA[85]) activates expression of *tcpPH* independent of ToxR (Supplementary Figure S4A). AphB is activated by a thiol-based switch mechanism, where oxygen-dependent modification of a single cysteine reside enhances its dimerization and transcriptional activation activity[86]. While *aphB* transcription is not significantly impacted by anaerobicity, its anaerobic activation and subsequent targets of transcriptional activation have been well-characterized[84]. Interestingly, *cadABC* is activated by AphB, providing a potential mechanistic link between the anaerobic bile condition and upregulation of *cadABC*. The vast majority of characterized AphB targets (13/18) were transcriptionally upregulated in at least two of the three overnight anaerobic bile comparisons in our RNA-seq dataset, further suggesting that AphB has robust activity in these conditions, This could help explain why we observed such robust activation of *cadABC* in conditions with relatively neutral pH. While CadC may not have been activated directly by the anaerobic bile condition, the cells could have achieved the robust activation of *cadABC* and thus maximize O1-antigen biosynthesis via post-translational activation of AphB. We hypothesize that an *aphB* mutant *V. cholerae* strain may display a defect in O1-antigen production in anaerobic bile conditions similar to the *cadC* and *cadA* mutant strains. This provides an indirect and ToxR-independent route for O1-antigen reduction to occur.

Overall, this RNA-seq dataset contributes novel confirmation and supporting evidence to many observations across the *V. cholerae* field over the last 30 years, highlighting the diversity of discoveries that have been made about the remarkable regulation governing *V. cholerae* environmental response and adaptation.
